# Tumor-specific neutrophils originating from meninges promote glioblastoma

**DOI:** 10.1101/2023.05.23.542010

**Authors:** Jiarui Zhao, Di Wu, Jiaqi Liu, Yang Zhang, Penghui Cao, Chunzhao Li, Shixuan Wu, Mengyuan Li, Yiwen Cui, Yiwen Sun, Ence Yang, Nan Ji, Jing Yang, Jian Chen

**Affiliations:** Department of Neurobiology, School of Basic Medical Sciences, Capital Medical University, Beijing 100069, China; Chinese Institute for Brain Research, Beijing, Research Unit of Medical Neurobiology, Chinese Academy of Medical Sciences, Beijing 102206, China; State Key Laboratory of Membrane Biology, School of Life Sciences, Peking University, Beijing 100871, China; Changping Laboratory, Beijing 102206, China; Peking University-Tsinghua University-National Institute Biological Sciences Graduate Program, Academy for Advanced Interdisciplinary Studies, Peking University, Beijing 100871, China; Center for Life Sciences, Academy for Advanced Interdisciplinary Studies, Peking University, Beijing 100871, China; IDG/McGovern Institute for Brain Research, Peking University, Beijing 100871, China; Institute of Molecular Physiology, Shenzhen Bay Laboratory, Shenzhen 518055, Guangdong, China; Department of Neurosurgery, Beijing Tiantan Hospital, Capital Medical University, Beijing 100050, China; China National Clinical Research Center for Neurological Diseases, Beijing Tiantan Hospital, Capital Medical University, Beijing 100070, China; Department of Medical Bioinformatics, School of Basic Medical Sciences, Peking University Health Science Center, Beijing 100191, China

**Author notes:** Corresponding Authors (N.J.), (J.Y.), (J.C.). Authors contributed equally (Jiarui Zhao, Di Wu, Jiaqi Liu, Yang Zhang).

**Keywords:** Tumor-specific neutrophils, tumor immune microenvironment, glioma, glioblastoma, meninges

## Abstract

Glioma is one of the most aggressive human cancers with limited therapeutic options. Though research has extensively examined immune components in those malignant tumors, the pathophysiological mechanism establishing their immunosuppressive microenvironment remains incompletely characterized. In this study, we report for the first time the unique presence of tumor-specific neutrophils (TSNs) in human glioblastoma (GBM) tumors. This newly defined neutrophil subtype exhibits the high expression of several immunosuppressive genes (e.g., CD274 and IDO1) and is strongly correlated with glioma grades and poor prognosis of patients. TSNs with comparable gene signatures are similarly present in the tumors but not bone marrow or spleen of mouse glioma models. Blockage of TSN recruitment by either *Cxcl1*-knockout in glioma cells or *Cxcr2*-deletion in host mice significantly enhances antitumor immunity and inhibits tumor progression. Surprisingly, we further identify the meninges as the key extratumoral source of generating TSNs in both human GBM patients and mouse glioma models. These results have elucidated a novel mechanism of neutrophils designating the tumor immune microenvironment and the essential link of meningeal immunity to glioma.

## Main Text

The tumor immune microenvironment has a determinant role in cancers^1^. Indeed, CD8^+^ T cells and NK cells, the body’s central components of antitumor immunity, are subjected to diverse immunosuppressive mechanisms in tumors, which together lead to their evasion and metastasis^2^. For instance, it has been recognized that myeloid cells, particularly tumor-associated macrophages (TAMs), can establish such an immunosuppressive microenvironment through their expression of various immune-checkpoint molecules (e.g., PD-L1) and inhibitory factors (e.g., IL-10 and ARG1)^3^.

Glioma is one of the most aggressive human cancers with a poor prognosis^4^. Due to the heterogeneity of tumor cells, the blood-brain barrier, and the immunosuppressive microenvironment, glioma tumors are often resistant to currently available chemotherapies, targeted therapies, or immunotherapies^5,6^. Accordingly, decades of research have been pursued to unravel glioma tumorigenesis in the hope of developing more effective treatments against this deadly cancer. Of importance, it has long been documented that myeloid cells are abundantly present within glioma tumors^7,8^. Studies on TAMs or microglia have suggested the involvement of those immunosuppressive myeloid cells in regulating glioma onset and progression. Studies on TAMs or microglia have suggested the involvement of those immunosuppressive myeloid cells in regulating glioma onset and progression^9–12^. However, strategies targeting TAMs or their immunosuppressive signals produce only marginal effects on tumor growth or overall survival in clinical contexts^5,13^. Such observations have raised the possibility that unknown immune component(s) exist to control the disease process of glioma. Notably, recent studies on several peripheral cancers, e.g., lung cancer^14,15^, liver cancer^16^, and prostate cancer^17^, have reported the immunomodulatory functions of neutrophils in those disease contexts. Whether neutrophils may designate the immune microenvironment of glioma remains unclear.

### Tumor-specific neutrophils in human glioblastoma tumors and their correlation with prognosis

As the entry point of the study, we systematically profiled myeloid cells in human glioblastoma (GBM) tumors to identify novel components designating the immune microenvironment. CD45^+^ CD11b^+^ CD14^+^ and CD45^+^ CD11b^+^ CD15^+^ myeloid cells were enriched for single-cell RNA sequencing (scRNA-seq) analyses (Fig. 1A). After quality control, a total of 29,072 cells from the tumors of four GBM patients were obtained (Table S1). Consistent with previous reports on GBM tumors^7,8^, macrophages were abundantly identified (Fig. 1B and Supplemental Fig. 1A). Four macrophage subtypes could be defined, i.e., classical macrophages (Ma_LYZ), macrophages with the high expression of M2 marker *MRC1* (Ma_MRC1), macrophages with increased levels of angiogenesis-associated markers *ADAM8* and *BNIP3* (Ma_BNIP3), and macrophages with the cell proliferation marker *KI67* (Ma_MKI67). In addition, a microglial population was detected in human GBM tumors (Fig. 1B).

**Figure 1.**
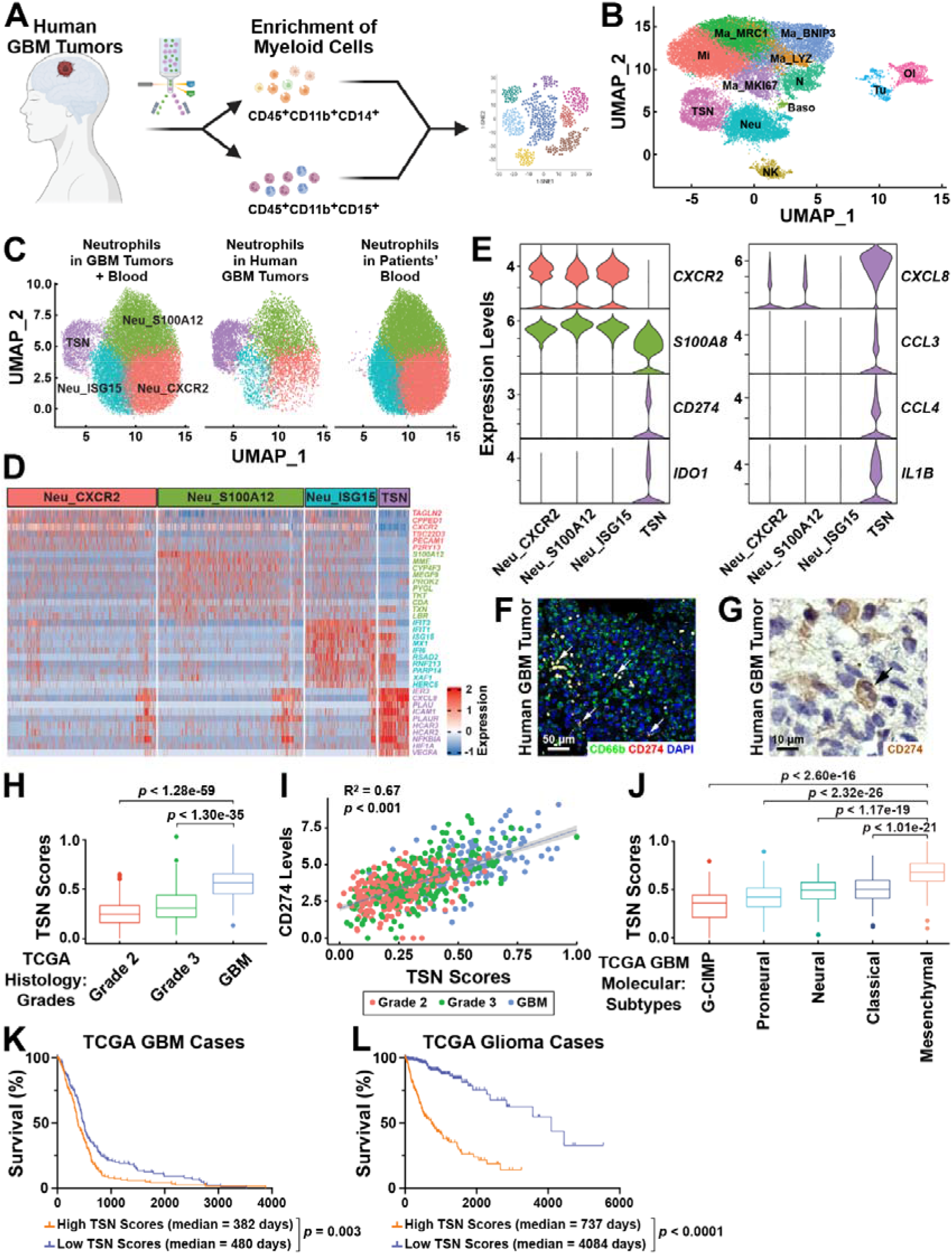
Tumor-specific neutrophils in human glioblastomas and their correlation with prognosis. **(A to G)** Identification of tumor-specific neutrophils (TSNs) in human glioblastomas (GBM). **(A)** Diagram of profiling myeloid cells in human GBM tumors. **(B)** Major cell types identified in human GBM tumors. Baso, basophils; Ma_MRC1, Ma_BNIP3, Ma_LYZ, Ma_MKI67, macrophages with corresponding markers; Mi, microglia; N, neurons; Neu, neutrophils; NK, natural killer cells; Ol, oligodendrocytes; TSN, tumor-specific neutrophils; Tu, tumor cells. **(C)** Neutrophil subtypes are defined in human GBM tumors and patients’ blood. Neu_CXCR2, Neu_ISG15, Neu_S100A12, neutrophils with corresponding markers; TSN, tumor-specific neutrophils. **(D)** Top expressing genes in neutrophil subtypes as defined in **(C)**. **(E)** Violin plot of the expression of neutrophil marker genes (*CXCR2* and *S100A8*), immunosuppressive genes (*CD274* and *IDO1*), and myeloid recruitment-related genes (*CXCL8*, *CCL3*, *CCL4*, and *IL1B*) in neutrophil subtypes. **(F and G)** Representative images of CD66b/CD274 immunofluorescence staining **(F)** or CD274 immunohistochemistry **(G)** of human GBM tumors. White arrows, CD66b^+^CD274^+^ cells; black arrow, CD274^+^ neutrophil with the lobulated nucleus. **(H to L)** TSNs are correlated with glioma grades and poor prognosis. **(H)** TSN scores are positively correlated with glioma grades in the TCGA dataset (Student’s *t*-test). **(I)** Correlation between TSN scores and CD274 expression levels in the TCGA dataset. The correlation coefficient (R) and *p*-value were calculated by Pearson correlation. **(J)** TSN scores are enriched in the mesenchymal subtype than in other molecular subtypes of human GBM (Student’s *t*-test). **(K and L)** Overall survival of TCGA GBM cases **(K)** or TCGA glioma cases **(L)** stratified by TSN scores (log-rank test).

Our profiling of myeloid cells in human GBM tumors revealed the presence of neutrophils (Fig. 1B). We also isolated neutrophils from GBM patients’ blood for comparison (Table S1). The combined analyses of neutrophils in human GBM tumors (6,503 neutrophils) and patients’ blood (21,329 neutrophils) could classify four subtypes (Fig. 1C to 1E), including neutrophils with the high expression of classical marker *CXCR2* (Neu_CXCR2), neutrophils with increased levels of maturation markers such as *MME* (Neu_S100A12), and neutrophils with the enrichment of interferon-stimulated genes (Neu_ISG15). Of importance, we observed the fourth subtype of neutrophils, which uniquely exhibited high levels of key immunosuppressive genes (Fig. 1E and Supplemental Fig. 1B), e.g., *CD274*, *CEACAM1*, *IDO1*, and *IDO2*, only in GBM tumors but not in patients’ blood (Fig. 1C and Supplemental Fig. 1C), and hence named as tumor-specific neutrophils (TSNs). In addition to the immunosuppressive genes, TSNs enriched the expression of myeloid recruitment-related genes (Fig. 1D and 1E), including *CXCL8*, *IL1B*, *CCL3*, and *CCL4*, suggesting their critical role in establishing the tumor immune microenvironment. Interestingly, TSNs showed decreased levels of *CXCR2* (Fig. 1E and Supplemental Fig. 1B), implicating their lower cell motility compared to other neutrophil subtypes^18^.

We confirmed the presence of TSNs, which were labeled as CD66b^+^CD274^+^ cells by immunofluorescence staining, in both human GBM and low-grade glioma tumors (Fig. 1F and Supplemental Fig. 1D). Indeed, the vast majority of CD274^+^ cells in GBM tumors overlapped with the neutrophil marker CD66b. Also, CD274^+^ cells with lobulated nuclei, the distinct morphological feature of neutrophils, were detected in human GBM tumors by immunohistochemistry (Fig. 1G).

To determine the clinical relevance of TSNs, a list of TSN signature genes (Table S2) was utilized to calculate “TSN scores” for the published databases of human gliomas (see **Materials & Methods**). In the TCGA dataset, which includes 513 GBM microarray data and 617 RNA-seq data of different glioma grades with clinical information^19,20^, TSN scores showed a positive correlation with glioma grades (Fig. 1H) or CD274 expression levels of tumors (Fig. 1I). In addition, TSN scores appeared highest in the mesenchymal subtype of GBM (Fig. 1J), which had the poorest prognosis among different molecular subtypes of GBM in the TCGA dataset. Accordingly, GBM patients (Fig. 1K) or glioma patients (Fig. 1L) with higher TSN scores were associated with lower overall survival rates. In parallel, the positive correlation between TSN scores and glioma grades was similarly observed in the CGGA dataset (Supplemental Fig. 2A), which contains RNA-seq data from 388 GBM and a total of 622 glioma samples^21^. Also, higher TSN scores of GBM patients (Supplemental Fig. 2B) or glioma patients (Supplemental Fig. 2C) were associated with poorer overall survival in the CGGA dataset.

To further corroborate this new identification of TSNs, we looked into the GBmap database comprising 26 published datasets that together contain approximately 1.1 million cells from the tumors of 240 glioma patients^22^. Within the GBmap dataset, 4,290 neutrophils were identified (Supplemental Fig. 3A). Notably, 86.6% of those neutrophils came from a single sample (Table S3), likely reflecting the technical challenges of profiling neutrophils due to their short lifespan and low RNA contents^16^. Nevertheless, neutrophils in the GBmap dataset and those in our current study could be clustered (Supplemental Fig. 3B), with a portion of neutrophils in the GBmap dataset defined as TSNs by their high TSN scores (Supplemental Fig. 3C).

### Tumor-specific neutrophils promote tumor progression in mouse glioma models

We next examined the presence of TSNs in mouse allograft glioma models. LCPN glioma cells and their derivative LCPNS-SIIN cells were generated (see **Materials & Methods**), which could effectively produce intracranial tumors with key features of gliomas in C57BL/6 wild-type mice (Supplemental Fig. 4). We profiled neutrophils from mouse LCPN tumors (Fig. 2A), as well as neutrophils from the spleens of control or tumor-bearing mice for comparison. A total of 92,181 cells, including 78,764 neutrophils, were obtained for analyses (Supplemental Fig. 5A). Six neutrophil subtypes were identified (Fig. 2B to 2D), i.e., immature neutrophils with the expression of secondary granule genes such as *Camp*, *Ltf*, and *Ngp* (Neu_Camp), neutrophils with increased levels of maturation markers (Neu_S100a6), neutrophils with the high expression of classical marker *Cxcr2* (Neu_Cxcr2), neutrophils with granulopoiesis and neutrophil extracellular traps (NETs) related genes *Slpi* (Neu_Slpi), neutrophils with the enrichment of interferon-stimulated genes (Neu_Isg15), and TSNs. Notably, the Neu_Camp subtype was almost absent in glioma tumors, and its population in the spleens decreased under the tumor-bearing condition (Fig. 2B), likely reflecting that Neu_Camp may have the ability to differentiate into other neutrophil subtypes (Supplemental Fig. 5B).

**Figure 2.**
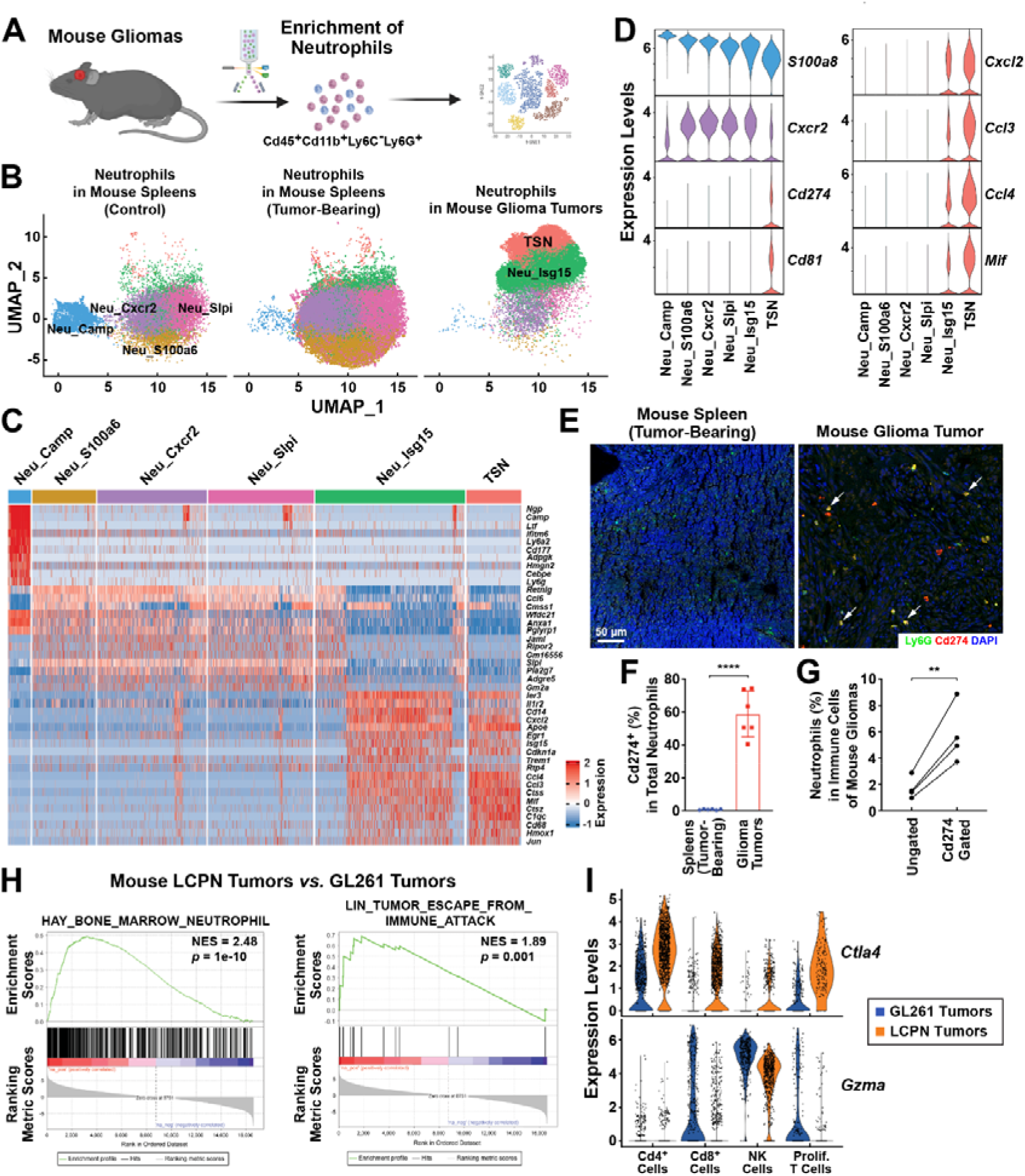
Tumor-specific neutrophils in mouse glioma models. C57BL/6 wild-type mice were intracranially implanted with glioma cell lines to establish allograft models. **(A to G)** Characterization of TSNs in mouse glioma models. **(A)** Diagram of profiling neutrophils in mouse gliomas. **(B)** TSNs and other neutrophil subtypes identified in mouse glioma tumors and spleens. Neu_Camp, Neu_Cxcr2, Neu_Isg15, Neu_S100a6, Neu_Slpi, neutrophils with corresponding markers; TSN, tumor-specific neutrophils. **(C)** Top expressing genes in neutrophil subtypes as defined in **(B)**. **(D)** Violin plot of the expression of neutrophil marker genes (*Cxcr2* and *S100a8*), immunosuppressive genes (*Cd274* and *Cd81*), and myeloid recruitment-related genes (*Ccl2*, *Ccl3*, *Ccl4*, and *Mif*) in neutrophil subtypes. **(E)** Representative images of Ly6G/Cd274 immunofluorescence staining of mouse spleen (tumor-bearing condition, left panel) and glioma tumor (right panel). **(F)** Percentage of TSNs (Cd45^+^ Cd274^+^ Cd11b^+^ Ly6G^+^) in total neutrophils (Cd45^+^ Cd11b^+^ Ly6G^+^) of the mouse spleens (tumor-bearing condition) and glioma tumors were quantified by FACS analyses (Student’s *t*-test). **(G)** Percentage of neutrophils (Cd45^+^ Cd11b^+^ Ly6G^+^) in total immune cells (Cd45^+^) of mouse glioma tumors without or with Cd274^+^ gating (Student’s *t*-test). ** *p* < 0.01, **** *p* < 0.0001. **(H and I)** TSNs designate the immune microenvironment of mouse gliomas. **(H)** Gene set enrichment plot for the neutrophil and immune-evasion signatures of LCPN *vs.* GL261 glioma tumors. **(I)** Violin plot of the expression of immunosuppressive genes (*Ctla4* and *Gzma*) in Cd4^+^ T cells, Cd8^+^ T cells, NK cells, and proliferating T cells within the tumor microenvironment of GL261 *vs.* LCPN glioma tumors.

Mouse TSNs exhibited gene signatures comparable to those in human counterparts, i.e., the unique expression of immunosuppressive genes such as *Cd274* and *Cd81*, increased levels of myeloid recruitment-related genes *Cxcl2*, *Ccl3*, *Ccl4*, and *Mif*, and the reduction of classical marker *Cxcr2* (Fig. 2C and 2D). Additionally, consistent with the observation in human GBM tumors, mouse TSNs were predominantly detected in glioma tumors but not in the spleens (Fig. 2B and Supplemental Fig. 5C). We verified this restricted presence of TSNs, which were labeled as Ly6G^+^Cd274^+^ cells, in mouse glioma tumors by immunofluorescence staining (Fig. 2E). Also, FACS analyses showed that TSNs counted for over 50% of neutrophils in glioma tumors (Fig. 2F), and Cd274 gating could be utilized as an effective marker to enrich neutrophils from total immune cells of glioma tumors (Fig. 2G).

We noticed that mouse GL261 glioma cells, which are commonly used in the research field^23^, produced intracranial tumors with significantly lower levels of neutrophil recruitment than that occurring in LCPN tumors (Supplemental Fig. 6A). Importantly, a stronger neutrophil response of LCPN tumors correlated with their enrichment of the pathways of tumor immune evasion and T-cell apoptosis, as assessed by bulk RNA-seq analyses (Fig. 2H and Supplemental Fig. 6B). In addition, we profiled non-tumor cells from the microenvironment of LCPN *vs.* GL261 glioma tumors (Supplemental Fig. 6C). Although the overall patterns of immune cells appeared comparable between the two tumor types (Supplemental Fig. 6D and 6E), Cd8^+^ T cells and proliferating total T cells in LCPN tumors showed the enhanced expression of the negative regulator *Ctla4* but a decrease of the effector protein *Gzma* (Fig. 2I). These results implicated that TSNs may promote the immunosuppressive microenvironment of mouse gliomas.

We went on to investigate the function of TSNs in mouse glioma models. Expression profiling revealed that LCPN tumors had higher levels of the neutrophil recruitment-related chemokines *Cxcl1* and *Cxcl5* than those in GL261 tumors (Fig. 3A). This differential expression of chemokines was confirmed by quantitative PCR analyses of *in vitro* cultured GL261 and LCPNS-SIIN cells (Fig. 3B). Because *Cxcl1* appeared as the predominantly expressed neutrophil recruitment-related chemokine, we generated the genetic deletion of *Cxcl1* in LCPNS-SIIN cells using the CRISPR/Cas9 approach (Supplemental Fig. 7A and 7B). *In vitro* cultured LCPNS-SIIN-*Cxcl1*KO cells underwent the same rate of cell proliferation as that of LCPNS-SIIN “wild-type” cells (Supplemental Fig. 7C). However, the recruitment of neutrophils (Fig. 3C) or TSNs (Fig. 3D) was significantly diminished in intracranial LCPNS-SIIN-*Cxcl1*KO tumors compared to that in LCPN-SIIN control tumors. In support of the immunosuppressive function of TSNs, the infiltration of Cd8^+^ T cells was elevated in LCPNS-SIIN-*Cxcl1*KO tumors (Fig. 3E and 3F), and those Cd8^+^ T cells exhibited less exhaustion (Fig. 3G). As a result, tumor growth of LCPNS-SIIN-*Cxcl1*KO cells was inhibited in C57BL/6 wild-type mice (Supplemental Fig. 7D), and the mice survived remarkably longer to the extent that almost no mortality was observed (Fig. 3H). Further, consistent with the high expression of myeloid recruitment-related factors by TSNs, the accumulation of tumor-associated macrophages/monocytes was also reduced in LCPNS-SIIN-*Cxcl1*KO tumors (Supplemental Fig. 7F). Of importance, the survival of nude mice, which lacked adaptive immune cells, was extended when intracranially implanted with LCPNS-SIIN-*Cxcl1*KO cells compared to LCPNS-SIIN control cells (Supplemental Fig. 7E), supporting that TSNs can modulate other myeloid cells in shaping the tumor microenvironment.

**Figure 3.**
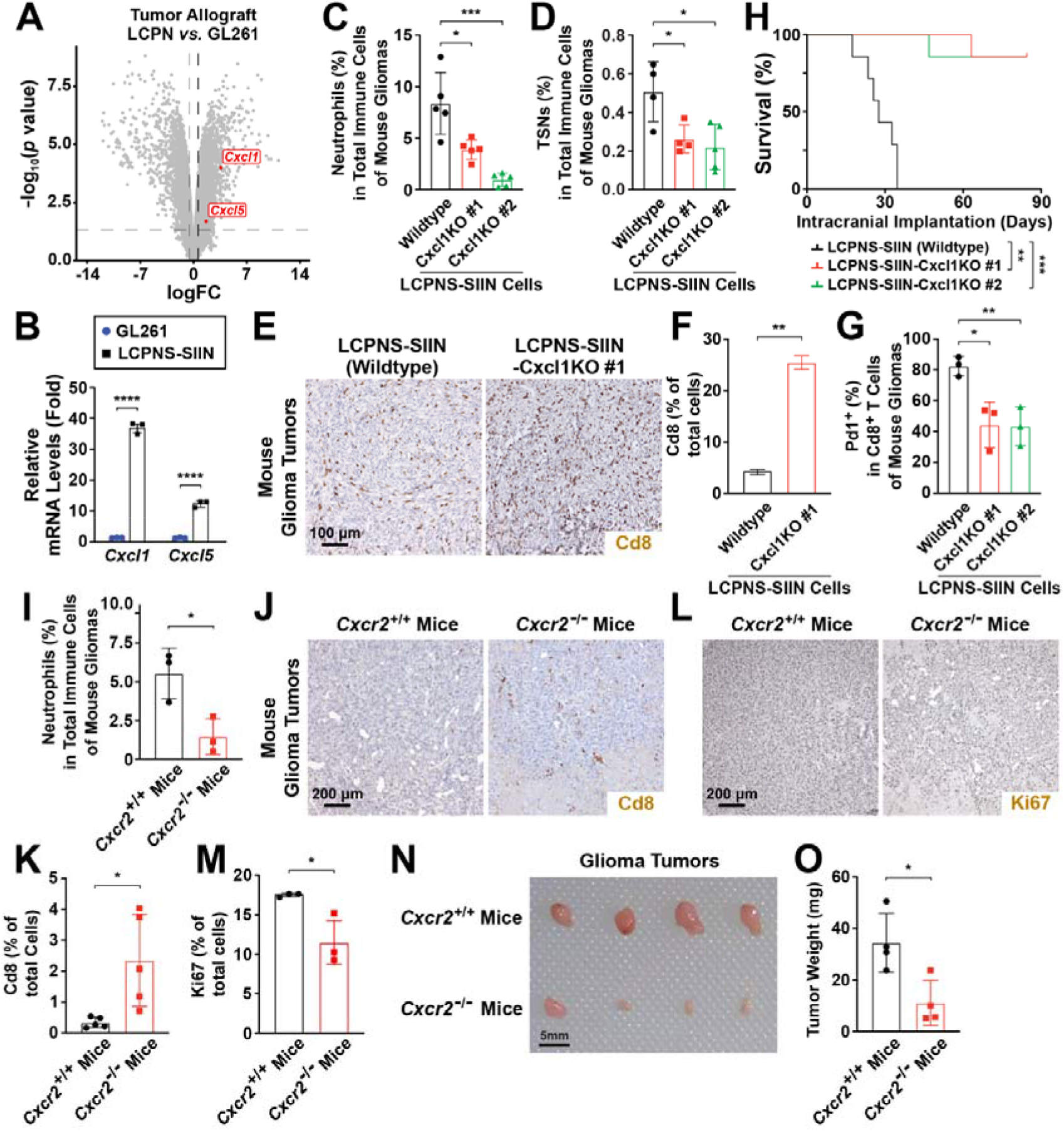
Neutrophils promote tumor progression in mouse glioma models. **(A and B)** Differential expression of neutrophil recruitment-related chemokines in mouse glioma models. **(A)** Volcano plot comparing the bulk RNA-seq data of LCPN *vs.* GL261 tumor allografts intracranially implanted in C57BL/6 wild-type mice. Increased expression levels of *Cxcl1* and *Cxcl5* in LCPN tumors are highlighted. **(B)** mRNA levels of *Cxcl1* and *Cxcl5* in cultured GL261 and LCPN-SIIN cells were assessed by quantitative PCR analyses (Student’s *t*-test). **(C to H)** Cxcl1 deletion in glioma cells inhibits TSN recruitment and tumor growth. C57BL/6 wild-type mice were intracranially implanted with LCPNS-SIIN cells or LCPNS-SIIN-Cxcl1KO cells (clone #1 or clone #2). **(C and D)** The percentage of neutrophils (Cd45^+^ Cd11b^+^ Ly6G^+^) **(C)** or TSNs (Cd45^+^ Cd274^+^ Cd11b^+^ Ly6G^+^) **(D)** in total immune cells (Cd45^+^) of tumors were examined by FACS analyses (Student’s *t*-test). **(E and F)** Representative images **(E)** and quantification **(F)** of Cd8 immunohistochemistry of mouse glioma tumors (Student’s *t*-test). **(G)** The percentage of Pd1^+^ population in Cd8^+^ T cells of glioma tumors was quantified by FACS analyses (Student’s *t*-test). (**H**) Survival curves of the mice (log-rank test). **(I to O)** Cxcr2 deletion in host mice blocks neutrophil recruitment and tumor progression. *Cxcr2 ^-/-^*or control *Cxcr2 ^+/+^*littermates were intracranially implanted with LCPNS-SIIN cells. **(I)** The percentage of neutrophils (Cd45^+^ Cd11b^+^ Ly6G^+^) in total immune cells (Cd45^+^) of tumors were examined by FACS analyses (Student’s *t*-test). **(J and K)** Representative images (**J**) and quantification **(K)** of Cd8 immunohistochemistry of mouse glioma tumors. **(L and M)** Representative images **(L)** and quantification **(M)** of Ki67 immunohistochemistry of glioma tumors. **(N and O)** Gross appearance **(N)** and tissue weights **(O)** of tumors in *Cxcr2 ^-/-^* or control *Cxcr2 ^+/+^* mice (Student’s *t*-test). * *p* < 0.05, ** *p* < 0.01, *** *p* < 0.001, **** *p* < 0.0001.

Conversely, we generated GL261-SIIN-*Cxcl1* cells stably overexpressing mouse Cxcl1 protein to boost their ability to recruit TSNs. As expected, TSNs became more abundant in intracranial GL261-SIIN-*Cxcl1* tumors than GL261-SIIN control tumors (Supplemental Fig. 8A). Also, the infiltration of Cd8^+^ T cells was markedly suppressed in GL261-SIIN-*Cxcl1* tumors (Supplemental Fig. 8B). Moreover, GL261-SIIN-*Cxcl1* glioma tumors became more aggressive, and the overall survival of the mice was worsened (Supplemental Fig. 8C).

Because Cxcl1 primarily acts via the specific receptor Cxcr2, we tested the effect of *Cxcr2* deletion in host mice on glioma progression. *Cxcr2^-/-^* or control *Cxcr2^+/+^* littermates were intracranially implanted with LCPNS-SIIN glioma cells. FACS analyses showed that TSN recruitment was strongly blocked in the tumors of *Cxcr2^-/-^* mice (Fig. 3I). Meanwhile, those tumors exhibited more infiltration of total T cells (Supplemental Fig. 8E and 8F) or Cd8^+^ T cells (Fig. 3J and 3K). Consistent with such enhanced antitumor immunity, *Cxcr2^-/-^* mice tumors had less cell proliferation than those in control *Cxcr2^+/+^* mice (Fig. 3L and 3M). Accordingly, tumor growth was evidently inhibited in *Cxcr2^-/-^* mice (Fig. 3N and 3O, and Supplemental Fig. 8D), with the overall survival of those mice being prolonged (Supplemental Fig. 8G). These results supported the immunosuppressive function of TSNs in promoting mouse glioma models.

### Myelopoiesis in the meninges generates tumor-specific neutrophils

We sought to determine the source of TSNs in mouse glioma models. FACS analyses did not detect any significant presence of TSNs in the bone marrow, blood, spleens, or cervical lymph nodes of mice intracranially implanted with glioma tumors (data not shown). We thus turned our attention to the meninges and profiled their cell populations. A diversity of immune cells, e.g., hematopoietic progenitor cells, lymphocytes, and myeloid cells, could be identified, with neutrophils being the most abundant cell type in mouse meninges (Supplemental Fig. 9A and 9B). We combined the analyses of neutrophils in mouse meninges (6,623 from control mice and 7,793 from tumor-bearing mice), spleens (15,093 from control mice and 37,376 from tumor-bearing mice), and glioma tumors (26,519). In addition to the six neutrophil subtypes as defined above (Fig. 2B and Supplemental Fig. 5B), the inclusion of meningeal neutrophils resulted in an additional subtype (Supplemental Fig. 10A and 10B), i.e., neutrophils with the cell proliferation marker *Ki67* (Neu_Mki67). Neu_Mki67 could represent progenitor cells that differentiated into other neutrophil subtypes (Supplemental Fig. 10C) and were predominantly enriched in the meninges of both control and tumor-bearing mice (Supplemental Fig. 10D).

Surprisingly, we discovered that the tumor-bearing condition robustly triggered the appearance of TSNs in mouse meninges (Fig. 4A and 4B), indicating this barrier tissue of the brain as the extratumoral source of TSNs. Such tumor-triggered meningeal TSNs were verified by FACS analyses, with the TSN percentage in total neutrophils being comparable between the meninges and glioma tumors (Fig. 4C). In addition, even before any detectable recruitment of neutrophils into tumors at five days after glioma cell implantation (Fig. 4D), granulocyte-monocyte progenitor (GMP) cells were already induced within the meninges (Fig. 4E), demonstrating the occurrence of meningeal myelopoiesis. Consistent with this notion, there was a significantly higher portion of proliferating neutrophils in the meninges than in the spleens or tumors of *Ki67-RFP* reporter mice after glioma implantation (Fig. 4F). In support of the meningeal contribution of neutrophils, those immune cells became accumulated in the meninges of *Cxcr2^-/-^* mice once their recruitment to glioma tumors was impaired (Fig. 4G).

**Figure 4.**
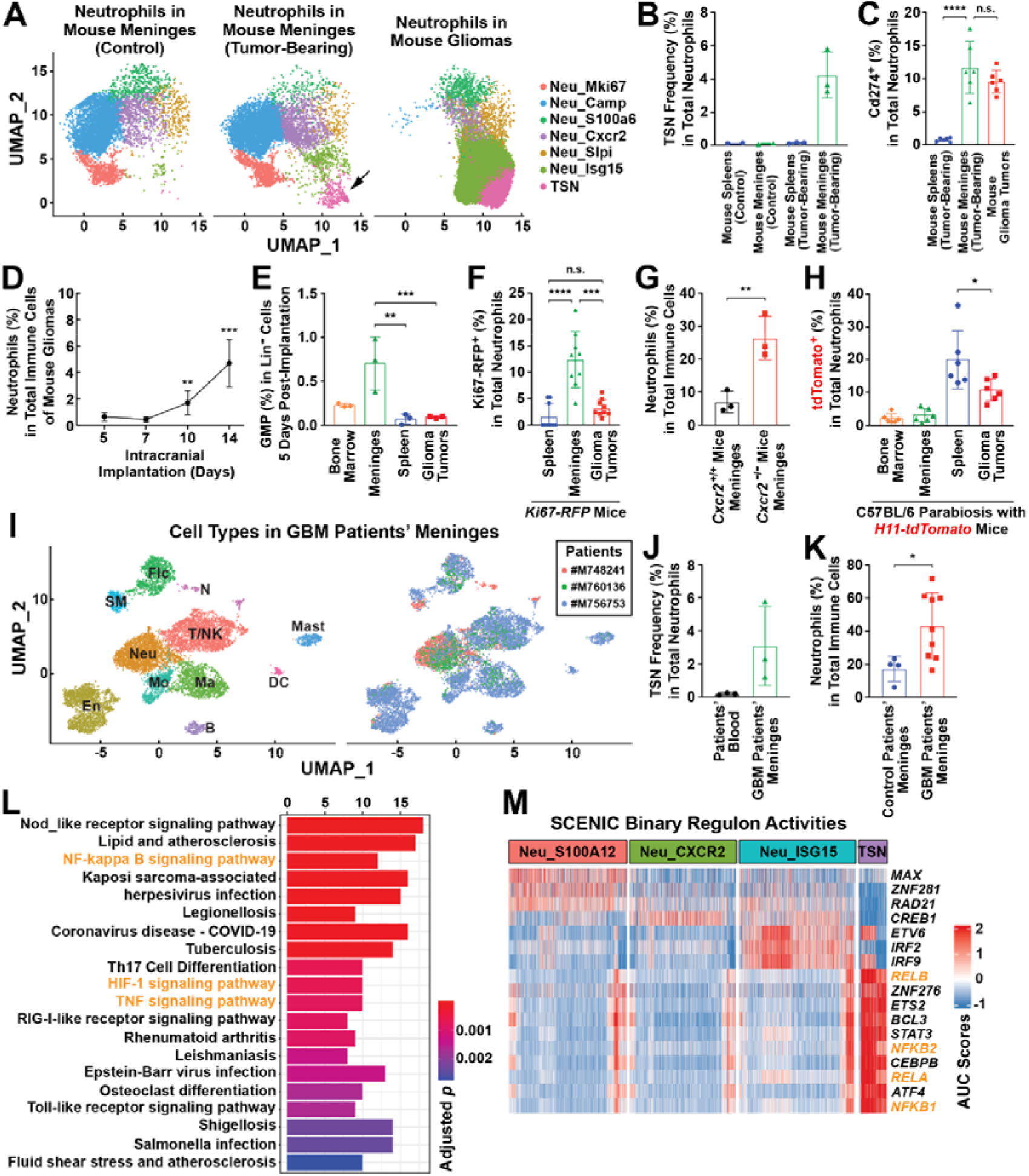
Myelopoiesis in the meninges generates tumor-specific neutrophils. **(A to C)** Presence of TSNs in the meninges of mouse glioma models. C57BL/6 wild-type mice were intracranially implanted with glioma cells. **(A)** TSNs and other neutrophil subtypes identified in mouse meninges and glioma tumors. Neu_Camp, Neu_Cxcr2, Neu_Isg15, Neu_Mki67, Neu_S100a6, Neu_Slpi, neutrophils with corresponding markers; TSN, tumor-specific neutrophils. The black arrow denotes TSNs in the mouse meninges of tumor-bearing condition. **(B)** TSN frequency in the total neutrophils of mouse spleens and meninges defined by scRNA-seq. **(C)** The percentage of TSNs (Cd45^+^ Cd274^+^ Cd11b^+^ Ly6G^+^) in neutrophils (Cd45^+^ Cd11b^+^ Ly6G^+^) of the spleens, meninges, and gliomas of tumor-bearing mice was quantified by FACS analyses (Student’s *t*-test). (**D to G**) Tumor-induced myelopoiesis in the meninges. C57BL/6 wild-type mice were intracranially implanted with glioma cells. **(D)** The percentage of neutrophils (Cd45^+^ Cd11b^+^ Ly6G^+^) in total immune cells (Cd45^+^) of tumors at different time points post-implantation. **(E)** The percentage of granulocyte-monocyte progenitors (GMP, Lin^-^ Sca-1^-^ c-Kit^+^ Cd34^+^ Cd16/32^+^; Lin: B220, Cd2, Cd3, Cd5, Cd8, Gr-1, and Ter119) in total Lin^-^ cells of the bone marrow, meninges, spleens, and gliomas of tumor-bearing mice were determined at day 5 post-implantation by FACS analyses (Student’s *t*-test). **(F)** Neutrophil proliferation occurs in the meninges. *Ki67-RFP* reporter mice were intracranially implanted with glioma cells. The percentage of Ki67-RFP^+^ cells in neutrophils (Cd45^+^ Cd11b^+^ Ly6G^+^) of the spleens, meninges, and tumors were assessed by FACS analyses (Student’s *t*-test). (**G**) Cxcr2 deletion in host mice retains neutrophils in the meninges. *Cxcr2 ^-/-^* or control *Cxcr2 ^+/+^* mice were intracranially implanted with glioma cells. The percentage of neutrophils (Cd45^+^ Cd11b^+^ Ly6G^+^) in total immune cells (Cd45^+^) of the meninges was quantified by FACS analyses (Student’s *t*-test). **(H)** Peripheral circulating immune cells were not sufficient to count for TSNs in mouse gliomas. A parabiosis between C57BL/6 wild-type mice and *H11-tdTomato* reporter mice was established. C57BL/6 mice in parabiotic pairs were then intracranially implanted with glioma cells. The percentage of tdTomato^+^ cells in neutrophils (Cd45^+^ Cd11b^+^ Ly6G^+^) of the bone marrow, meninges, spleens, and tumors of C57BL/6 mice was determined by FACS analyses (Student’s *t*-test). (**I to K**) Presence of TSNs in the meninges of human GBM patients. (**I**) Major cell types identified in GBM patients’ meninges. B, B cells; DC, dendritic cells; En, endothelial cells; Flc, fibroblast-like cells; Ma, macrophages; Mast, mast cells; Mo, monocytes; N, neurons; Neu, neutrophils; SM, smooth muscle cells; T/NK, T cells / natural killer cells. **(J)** TSN frequency in the total neutrophils of GBM patients’ blood and meninges defined by scRNA-seq. **(K)** The percentage of neutrophils (CD45^+^ CD11b^+^ CD14^-^ CD15^+^ HLA-DR^-^ CD33^low^ CD66b^+^) in total immune cells (CD45^+^) of the meninges of GBM or control patients was examined by FACS analyses (Student’s *t*-test). **(L and M)** Regulation of human TSN development by distinct signaling pathways. **(L)** KEGG enrichment diagram of human TSN marker genes. **(M)** Heat map of SCENIC binary regulon activities in TSNs and other neutrophil subtypes. Components of the NF-κB pathway are highlighted in orange. * *p* < 0.05, ** *p* < 0.01, *** *p* < 0.001, **** *p* < 0.0001.

To further corroborate the meningeal contribution of neutrophils to glioma tumors, we performed the parabiosis procedure (Supplemental Fig. 10E), which has been exploited in the research field to distinguish immune cells derived from peripheral circulation and the meninges^24,25^. There was an abundant presence (35% ∼ 45%) of tdTomato^+^ cells among total immune cells in the spleens and blood of C57BL/6 wild-type mice connected to *H11-tdTomato* reporter mice (Supplemental Fig. 10F), showing the successful establishment of parabiotic pairs. In contrast, chimerism rates of the bone marrow and meninges were low (< 5%) due to their regionalized hematopoiesis, as previously reported^25^. We intracranially implanted glioma cells into C57BL/6 wild-type mice in parabiosis with *H11-tdTomato* mice. Of importance, only 10% of neutrophils in glioma tumors were tdTomato^+^, which was significantly lower than the percentage of tdTomato^+^ neutrophils in the spleens or blood of C57BL/6 wild-type mice (Fig. 4H), indicating that peripheral circulation was not sufficient to count for the neutrophil recruitment in mouse gliomas.

We then determined whether a similar neutrophil response of the meninges would occur in human GBM patients. To this end, we profiled meningeal cells from three GBM patients (Table S1). A variety of immune cells were identified in GBM patients’ meninges (Fig. 4I and Supplemental Fig. 11A), though their detailed patterns varied from those in mouse meninges as described above (Supplemental Fig. 9A), e.g., the B-cell population in human meninges was much smaller than that in mice.

Regardless, neutrophils still constituted a major population among human meningeal cells. More importantly, TSNs could be detected in all three patients’ meninges (Fig. 4J). On the contrary, very few TSNs were present in those patients’ blood, confirming the specificity of TSN identification (Fig. 4J). Furthermore, this tumor-triggered neutrophil response was confirmed by FACS analyses of the meninges of GBM patients or non-GBM control patients (Fig. 4K).

We finally explored the signaling pathways of TSN generation. The combined analyses of human neutrophils obtained in the current study suggested that TSNs might originate from normal mature neutrophils (Supplemental Fig. 11B). Also, the NF-κB and HIF signaling pathways were significantly enriched in human TSNs (Fig. 4L). Indeed, SCENIC^26^ identified that several members of the NF-κB transcription factor family specifically controlled the genes expressed by human TSNs compared to other neutrophil subtypes (Fig. 4M). Accordingly, target genes of the NF-κB and HIF pathways were highly upregulated in human TSNs (Supplemental Fig. 11C), implicating that inflammatory and hypoxic cues may stimulate the TSN generation in GBM patients’ meninges and tumors.

In sum, this study has identified TSNs as a new, unique subtype of neutrophils present within human GBM tumors that are strongly correlated with glioma grades and poor prognosis of patients. It is worth noting that TSNs simultaneously express multiple immunosuppressive signals and myeloid recruitment-related factors, thus enabling their multifunctional involvement in shaping innate and adaptive immunity in the tumor microenvironment. Therefore, targeting any single aspect of TSNs, e.g., CD274-mediated inhibition of CD8^+^ T cells, may be insufficient to overcome their immunosuppressive arsenal. Instead, our current characterization of TSNs suggests the necessity to tackle them as an entity to achieve therapeutic benefits. Indeed, we have shown in mouse glioma models that blocking TSN recruitment can significantly inhibit the disease progression and even lead to the long-term survival of otherwise lethal tumors. How TSNs may collaborate with other myeloid cells, e.g., TAMs and microglia, in the immune microenvironment warrants more detailed investigations.

Recent studies on lung cancer^14,15^, liver cancer^16^, or prostate cancer^17^ have reported the immunomodulatory functions of neutrophils. In contrast to those previous reports, our work comparing neutrophil subtypes within glioma tumors *vs.* those in common immune compartments (e.g., spleen and blood) successfully defined TSNs with unique gene signatures, such as the enriched expression of CD274 and the downregulation of CXCR2. Of importance, this definition of TSNs may become applicable to other cancer types. For example, a similar TSN population could be identified in the neutrophils of mouse lung tumors by re-analyzing the published dataset^27^ (Supplemental Fig. 12), implicating a broad involvement of TSNs in the tumor immune microenvironment.

TSNs in human GBM and mouse glioma tumors exhibit a restricted presence and are largely absent in the bone marrow, spleen, lymph nodes, or blood. This distinct feature of TSNs may reflect the tissue location of glioma tumors inside the central nervous system. However, we have unexpectedly discovered the meninges, a barrier tissue of the brain, as the extratumoral source of generating TSNs. Though previous works have suggested the complex functions of meninges in hematopoiesis^28^, lymphopoiesis ^24^, and brain-specific antigen selection ^29^, the contribution of meningeal myelopoiesis to glioma had been uncharted until this study. The pathophysiological mechanism by which glioma tumors trigger TSN generation in the meninges demands future research efforts. Moreover, whether immunosuppressive neutrophil subtype(s) originating from the meninges might participate in other pathological conditions of the brain, e.g., autoimmune diseases or strokes, appears a tempting possibility to explore.

Together, we have elucidated a novel mechanism of neutrophils designating the tumor immune microenvironment and the essential link of meningeal immunity to glioma, which uncovers an exciting entry point for therapeutic applications.

## Acknowledgments

This work was funded by the National Key Research and Development Program of China (#2022YFA1103900 to J.C.; #2019YFA0802003 to J.Y.); the Changping Laboratory and the CAMS Innovation Fund for Medical Sciences (#2019-I2M-5-015 to J.C.); the National Natural Science Foundation of China (#31970974, #32061143007, #32125017, and #32150008 to J.Y.); the Beijing Natural Science Foundation (#7232086 to J.Y.). Additional funding for J.C. is from the Chinese Institute for Brain Research. J.Y. is also supported by the State Key Laboratory of Membrane Biology and the Center for Life Sciences at Peking University, and the Institute of Molecular Physiology at Shenzhen Bay Laboratory. The authors declare no conflicts of interest.

## Author Contributions

J.Z., W.D., J.L., P.C., S.W., M.L., Y.C., and Y.S. performed experiments. J.Z., J.L., Y.Z., C.L., and N.J. collected and processed human samples. J.Z., E.Y., N.J., Y.J., and J.C. analyzed the experimental data. J.Z., J.Y., and J.C. prepared the manuscript.

---

**Supplemental Figure 1.**
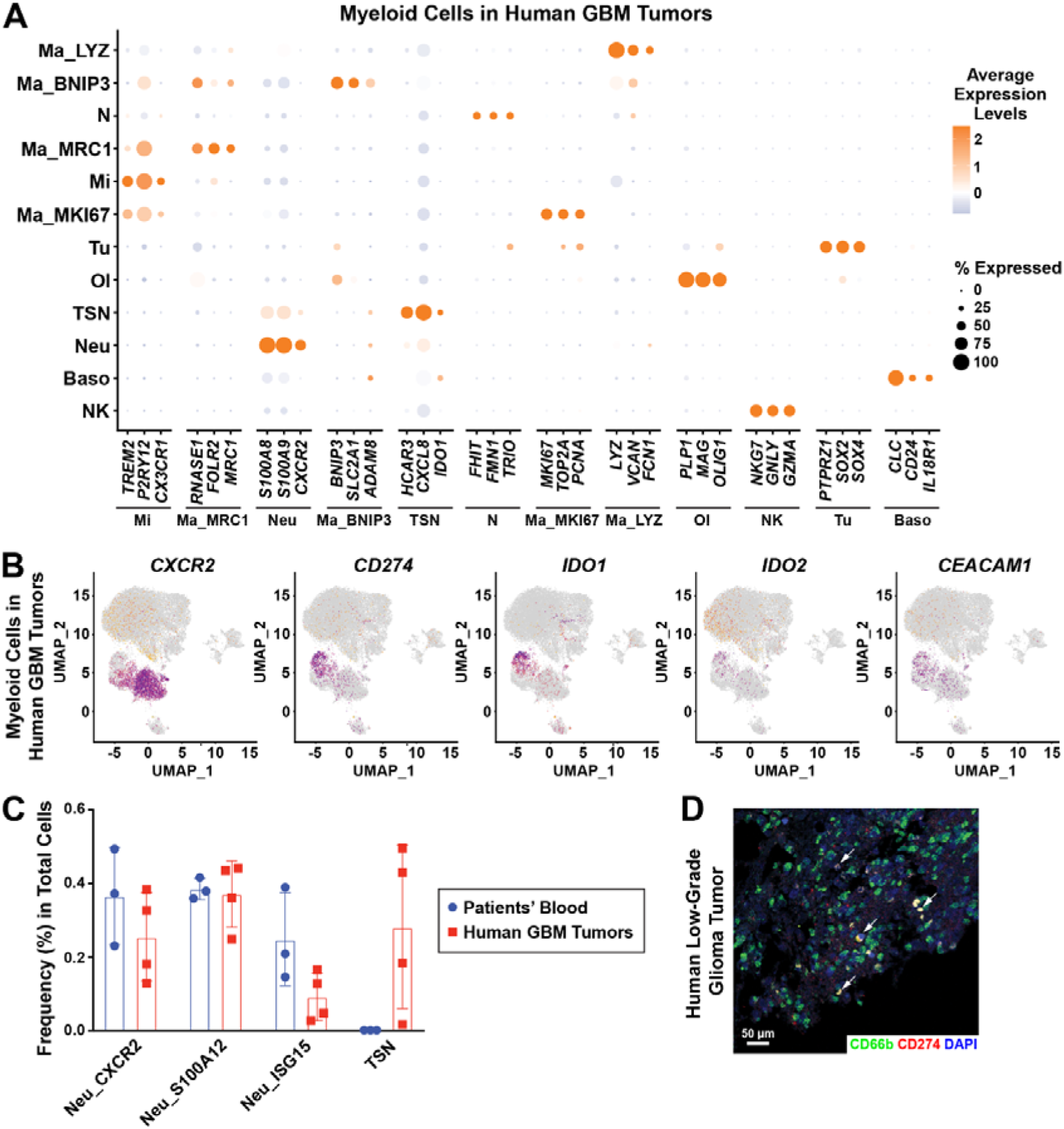
Characterization of neutrophil subtypes in human GBM tumors. Related to Figure 1. (A) Dot plot of major cell types identified by profiling myeloid cells in human GBM tumors. Three top marker genes for each cell type are shown. Baso, basophils; Ma_MRC1, Ma_BNIP3, Ma_LYZ, Ma_MKI67, macrophages with corresponding markers; Mi, microglia; N, neurons; Neu, neutrophils; NK, natural killer cells; Ol, oligodendrocytes; TSN, tumor-specific neutrophils; Tu, tumor cells. (B) Expression of the neutrophil marker gene (*CXCR2*) and immunosuppressive genes (*CD274*, *IDO1, IDO2*, and *CEACAM1*) in major cell types identified in **(A)**. (C) Frequency of TSNs and other subtypes in the total neutrophils of GBM patients’ blood and tumors defined by scRNA-seq. (D) Representative image of CD66b/CD274 immunofluorescence staining of human low-grade glioma tumor. White arrows, CD66b^+^CD274^+^ cells.

**Supplemental Figure 2.**
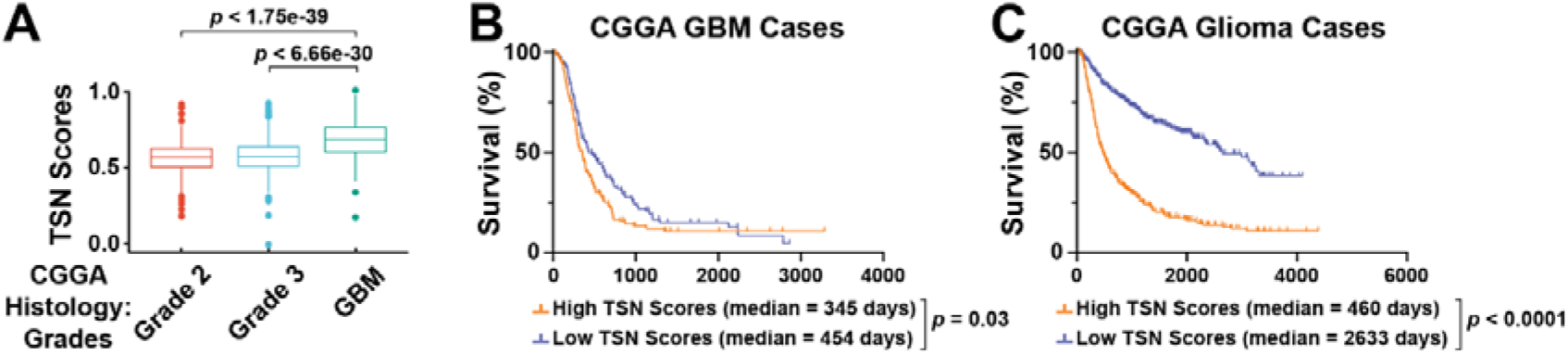
TSNs are correlated with glioma grades and poor prognosis. Related to Figure 1. **(A)** TSN scores are positively correlated with glioma grades in the CGGA dataset (Student’s *t*-test). **(B and C)** Overall survival of CGGA GBM cases **(A)** or CGGA glioma cases **(B)** stratified by TSN scores (log-rank test).

**Supplemental Figure 3.**
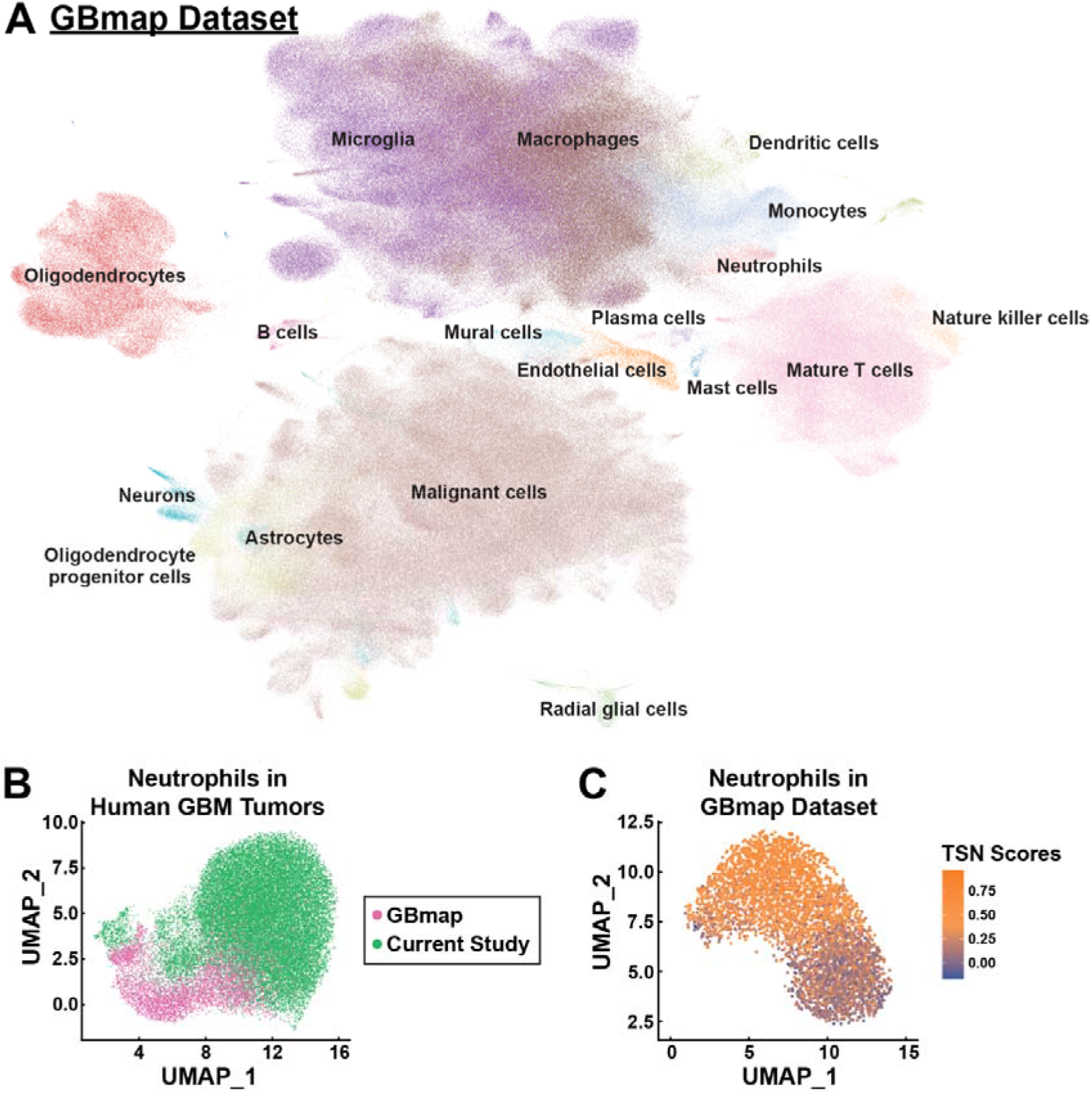
Neutrophils identified in the GBmap dataset. Related to Figure 1. **(A)** Major cell types of human gliomas identified in the GBmap dataset. **(B)** UMAP plot comparing the scRNA-seq results of neutrophils in the GBmap dataset and the current study. **(C)** TSN scores of neutrophils identified in the GBmap dataset.

**Supplemental Figure 4.**
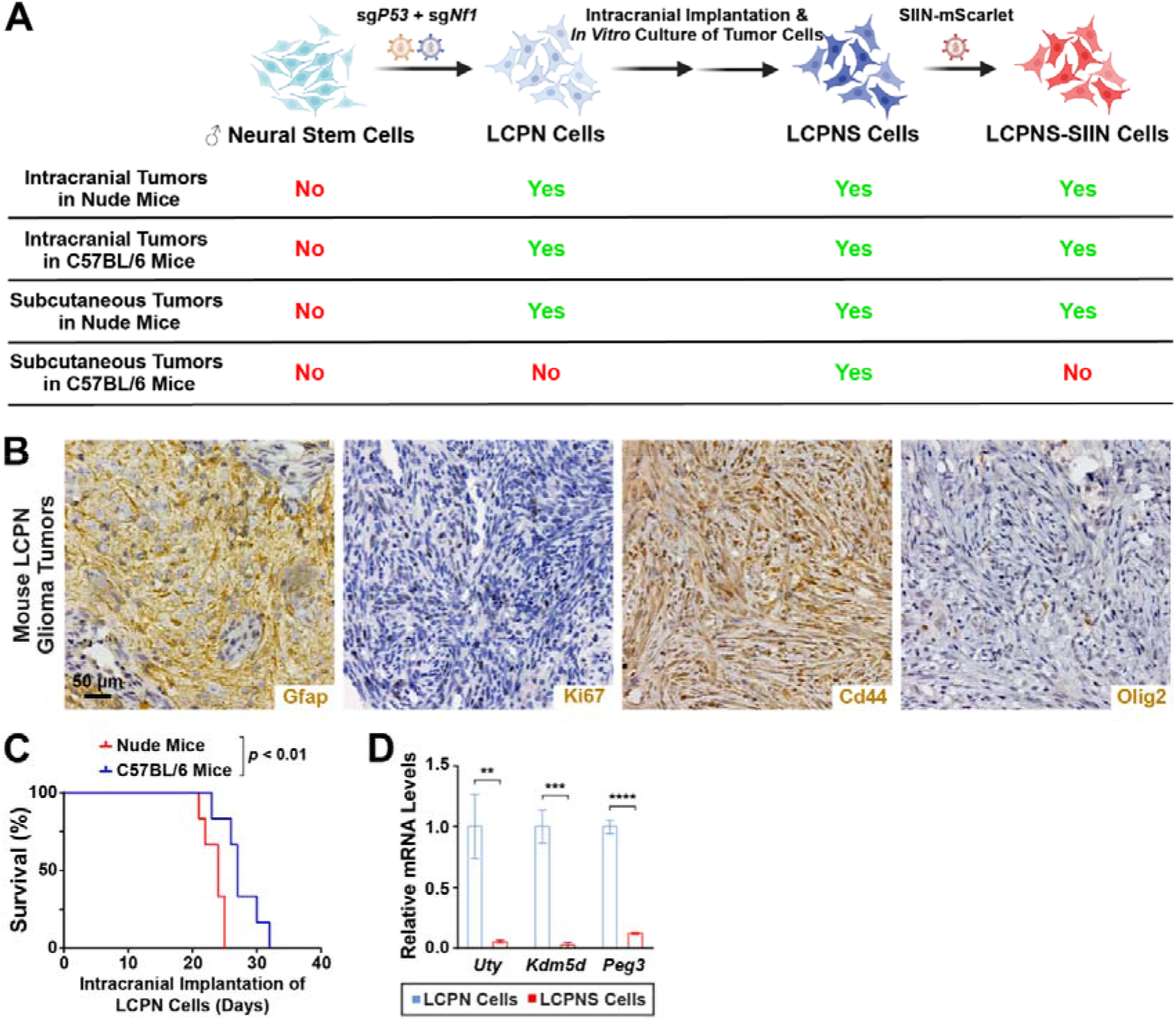
Characterization of mouse LCPN glioma cell lines. Related to Figure 2 and Figure 3. **(A)** Summary of the generation and tumorigenicity of mouse LCPN and derived cells. **(B)** Representative images of Gfap, Ki67, Cd44, or Olig2 immunohistochemistry of intracranial LCPN glioma tumors in C57BL/6 wild-type mice. **(C)** Survival curves of C57BL/6 wild-type or nude mice intracranially implanted with LCPN cells (log-rank test). **(D)** mRNA levels of Y-antigen genes (*Uty*, *Kdm5d*, and *Peg3*) in cultured LCPN and LCPNS cells were assessed by quantitative PCR analyses (Student’s *t*-test). ** *p* < 0.01, *** *p* < 0.001, **** *p* < 0.0001.

**Supplemental Figure 5.**
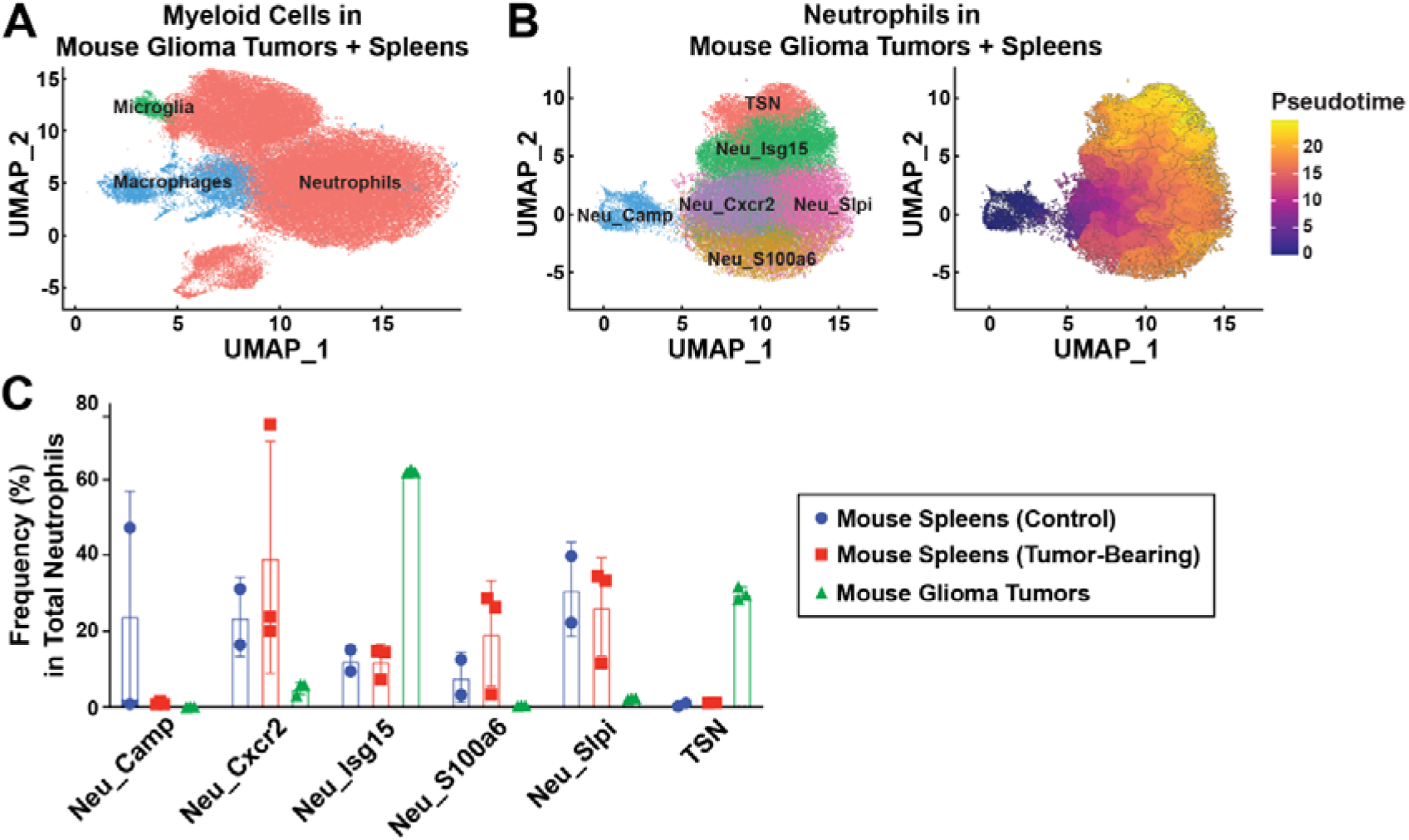
Characterization of TSNs in mouse glioma models. Related to Figure 2. **(A)** Major cell types identified by profiling neutrophils in mouse glioma tumors and the spleens of control or tumor-bearing mice. **(B)** UMAP plot (left panel) and pseudotime trajectory analysis (right panel) of TSNs and other neutrophil subtypes defined in mouse glioma tumors and the spleens of control or tumor-bearing mice. Neu_Camp, Neu_Cxcr2, Neu_Isg15, Neu_S100a6, Neu_Slpi, neutrophils with corresponding markers; TSN, tumor-specific neutrophils. **(C)** Frequency of TSNs and other subtypes in the total neutrophils of mouse glioma tumors and the spleens of control or tumor-bearing mice defined by scRNA-seq.

**Supplemental Figure 6.**
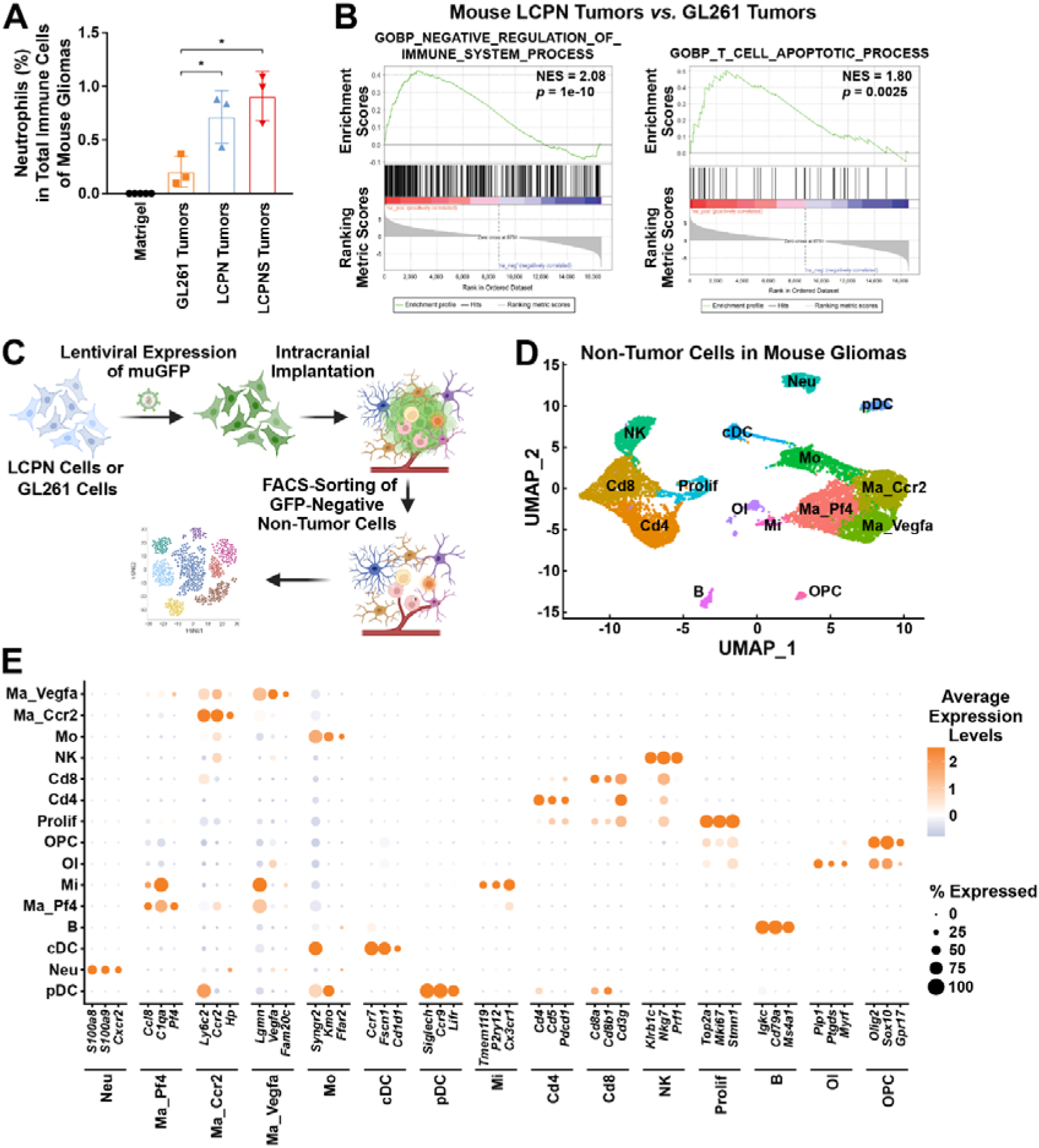
Comparison of mouse glioma models. Related to Figure 2 and Figure 3. **(A and B)** C57BL/6 wild-type mice were intracranially implanted with indicated glioma cells. **(A)** The percentage of neutrophils (Cd45^+^ Cd11b^+^ Ly6G^+^) in total immune cells (Cd45^+^) of the tumors of different glioma cells were examined by FACS analyses (Student’s *t*-test). * *p* < 0.05. **(B)** Enrichment plot for the gene sets negatively regulating immune processes or T cell apoptosis of LCPN *vs.* GL261 glioma tumors. (**C to E**) LCPN or GL261 cells stably transduced by muGFP-expressing lentiviruses were intracranially implanted into C57BL/6 wild-type mice, and GFP-negative non-tumor cells were sorted for scRNA-seq analyses **(C)**. **(D)** Major cell types identified in the microenvironment of LCPN or GL261 tumors. B, B cells; Cd4, Cd4^+^ T cells; Cd8, Cd8^+^ T cells; cDC, conventional dendritic cells; pDC, plasmacytoid dendritic cells; Ma_Ccr2, Ma_Pf4, Ma_Vegfa, macrophages with corresponding markers; Mi, microglia; Mo, monocytes; Neu, neutrophils; NK, natural killer cells; Ol, oligodendrocytes; OPC, oligodendrocyte progenitor cells; Prolif, proliferating T cells. **(E)** Dot plot of major cell types identified in **(D)**. Three top marker genes for each cell type are shown.

**Supplemental Figure 7.**
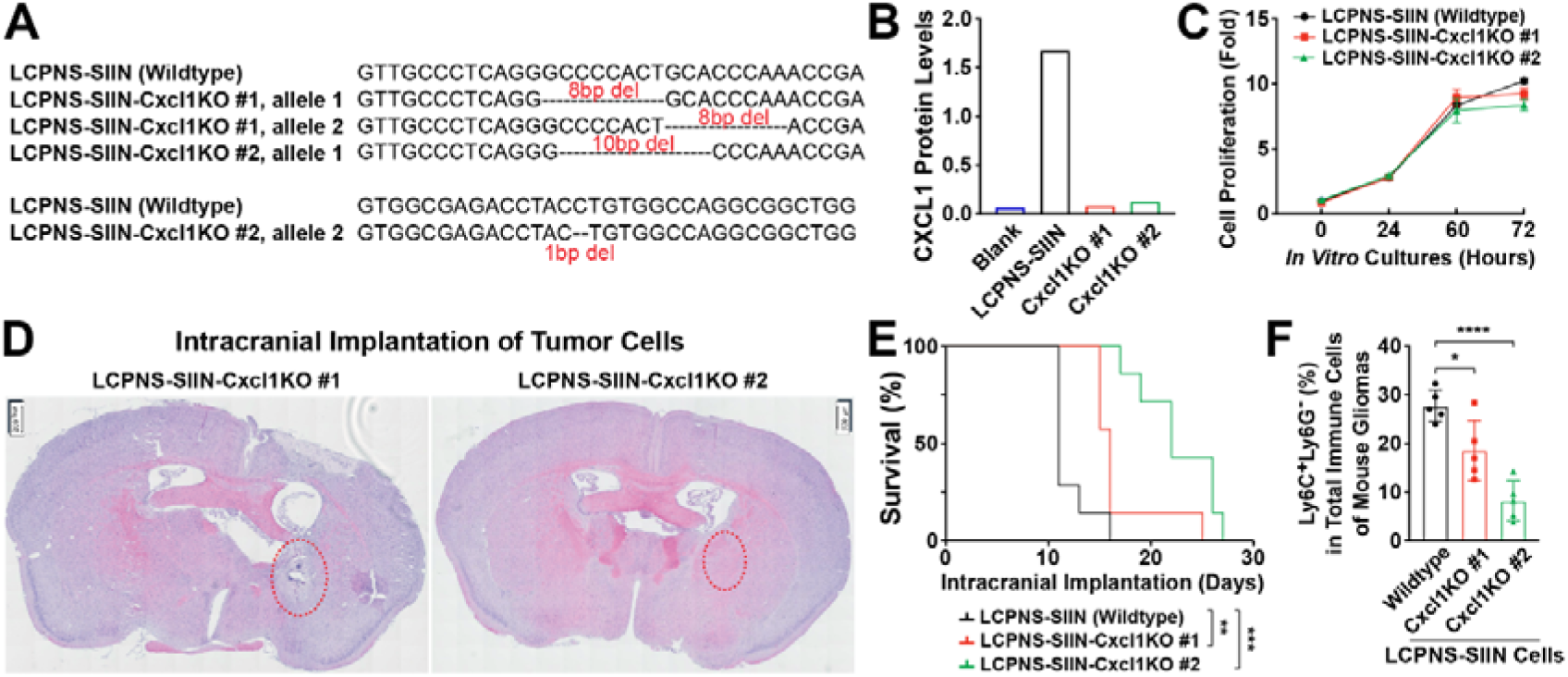
Cxcl1 deletion in glioma cells inhibits tumor growth. Related to Figure 3. **(A)** DNA sequencing results of the *Cxcl1* alleles of LCPNS-SIIN wild-type cells and LCPNS-SIIN-Cxcl1KO cells (clone #1 or clone #2). **(B)** Cxcl1 protein levels in the culture media of LCPNS-SIIN cells or LCPNS-SIIN-Cxcl1KO cells (clone # 1 or clone # 2) were quantified by ELISA analyses. **(C)** Proliferation rates of *in vitro* cultured LCPNS-SIIN cells or LCPNS-SIIN-Cxcl1KO cells (clone # 1 or clone # 2) were measured by CellTiter-Glo. **(D)** Representative H&E staining images of mouse brains implanted with LCPNS-SIIN-Cxcl1KO cells (clone # 1 or clone # 2). Red dashed lines denote the tumor sites. **(E)** Survival curves of nude mice intracranially implanted with LCPNS-SIIN cells or LCPNS-SIIN-Cxcl1KO cells (clone # 1 or clone # 2) (log-rank test). **(F)** C57BL/6 wild-type mice were intracranially implanted with LCPNS-SIIN cells or LCPNS-SIIN-Cxcl1KO cells (clone # 1 or clone # 2). The percentage of Cd11b^+^ Ly6C^+^ Ly6G^-^ cells in total immune cells (Cd45^+^) of tumors were examined by FACS analyses (Student’s *t*-test). * *p* < 0.05, ** *p* < 0.01, *** *p* < 0.001, **** *p* < 0.0001.

**Supplemental Figure 8.**
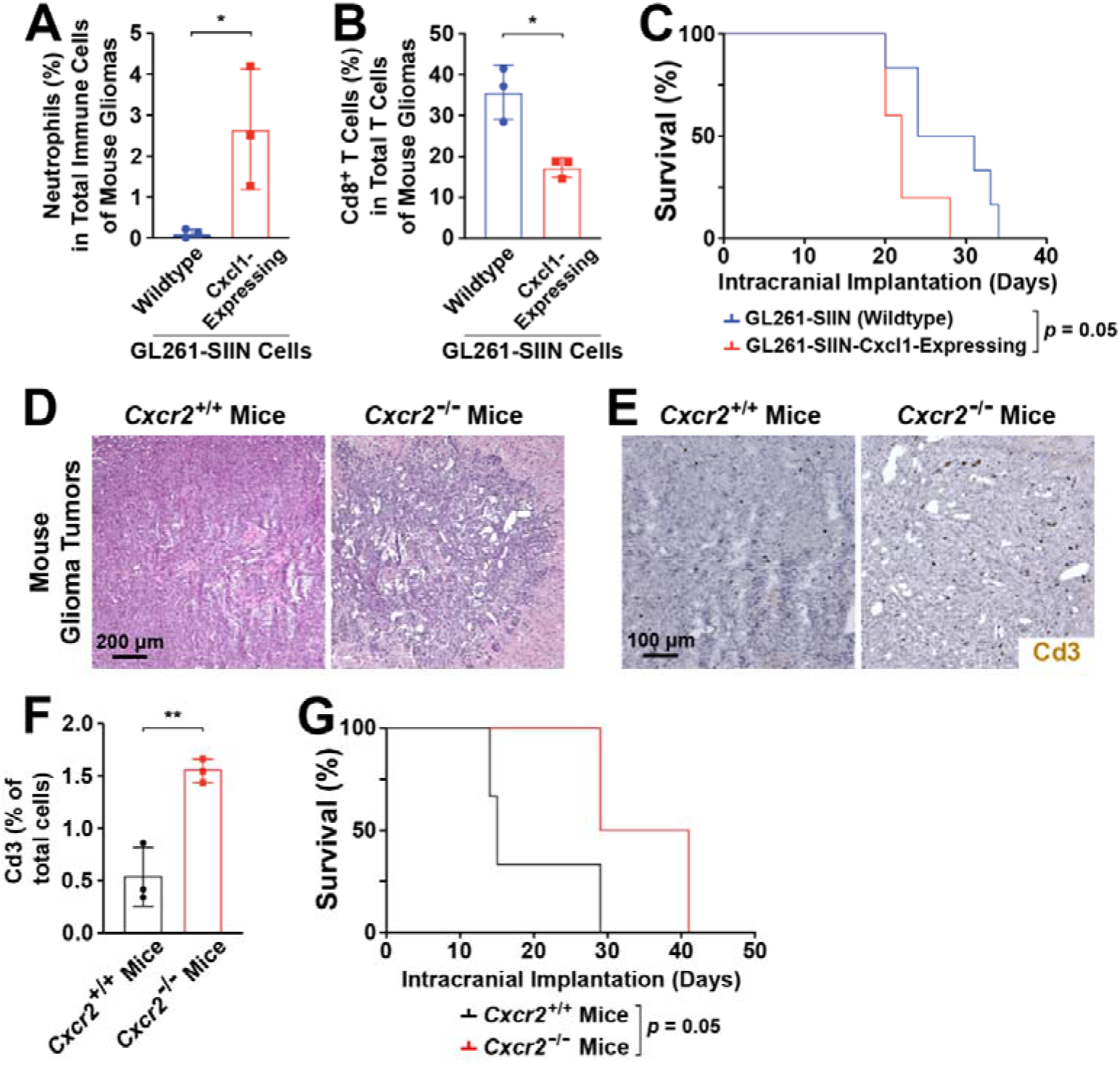
TSN recruitment promotes tumor growth in mouse glioma models. Related to Figure 3. **(A to C)** Cxcl1 overexpression in glioma cells promotes neutrophil recruitment and tumor growth. C57BL/6 wild-type mice were intracranially implanted with GL261-SIIN wild-type cells or GL261-SIIN-Cxcl1-expressing cells. (**A**) The percentage of neutrophils (Cd45^+^ Cd11b^+^ Ly6G^+^) in total immune cells (Cd45^+^) of tumors was examined by FACS analyses (Student’s *t*-test). (**B**) The percentage of Cd8^+^ cells in total T cells (Cd3^+^) of tumors was examined by FACS analyses (Student’s *t*-test). **(C)** Survival curves of the mice (log-rank test). **(D to G)** *Cxcr2 ^-/-^* or control *Cxcr2 ^+/+^* littermates were intracranially implanted with LCPNS-SIIN cells. **(D)** Representative H&E-staining images of tumors. **(E and F)** Representative images (**E**) and quantification **(F)** of Cd3 immunohistochemistry of mouse glioma tumors. (**G**) Survival curves of the mice (log-rank test). * *p* < 0.05, ** *p* < 0.01.

**Supplemental Figure 9.**
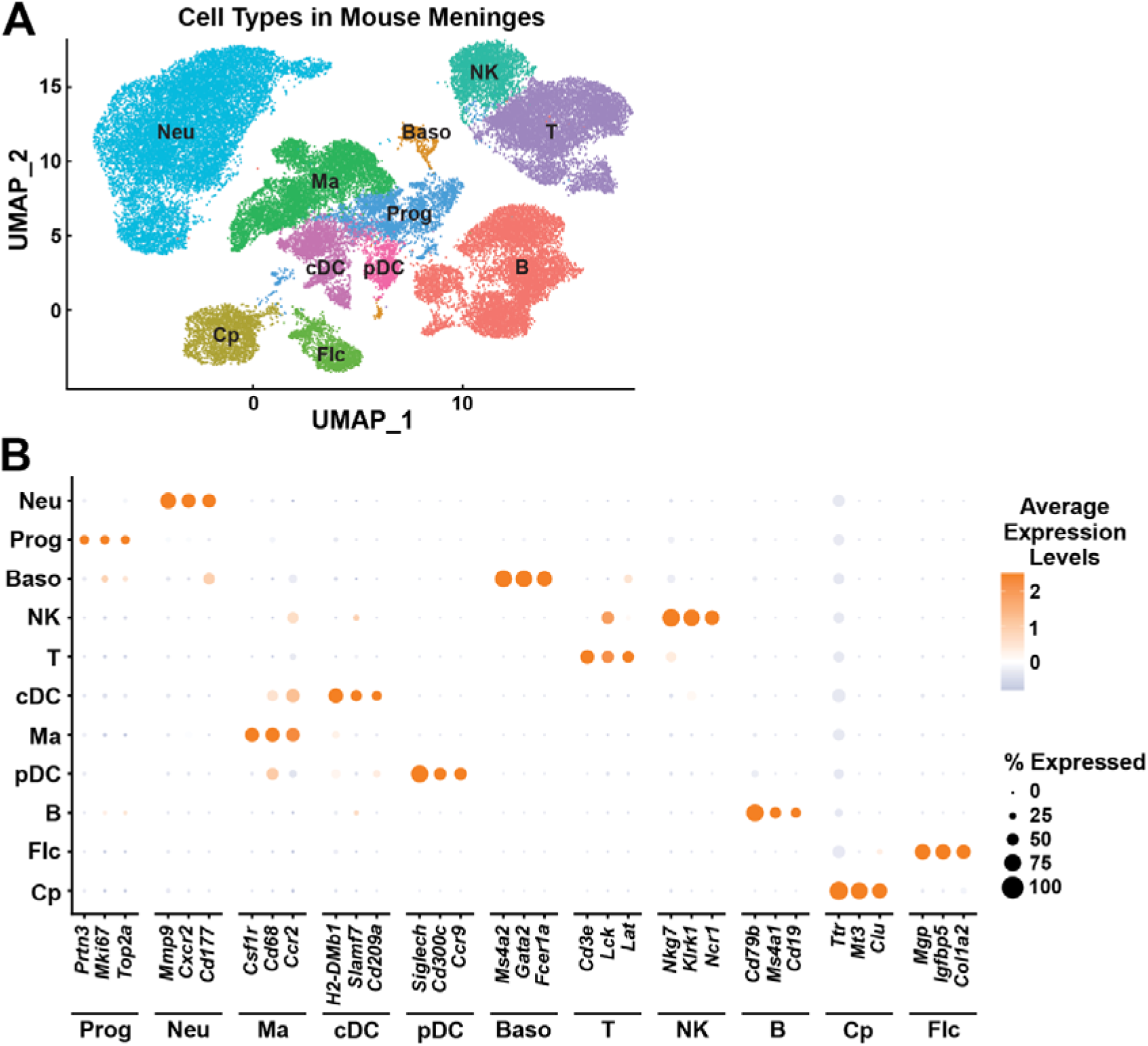
Characterization of meningeal cells in mouse glioma models. Related to Figure 4. C57BL/6 wild-type mice were intracranially implanted with glioma cells. **(A)** Major cell types identified in the meninges of control or tumor-bearing mice. B, B cells; Baso, basophils; Cp, choroid plexus cells; cDC, conventional dendritic cells; pDC, plasmacytoid dendritic cells; Flc, fibroblast-like cells; Ma, macrophages; Neu, neutrophils; NK, natural killer cells; Prog, hematopoietic progenitor cells; T, T cells. **(B)** Dot plot of major cell types identified in **(A)**. Three top marker genes for each cell type are shown.

**Supplemental Figure 10.**
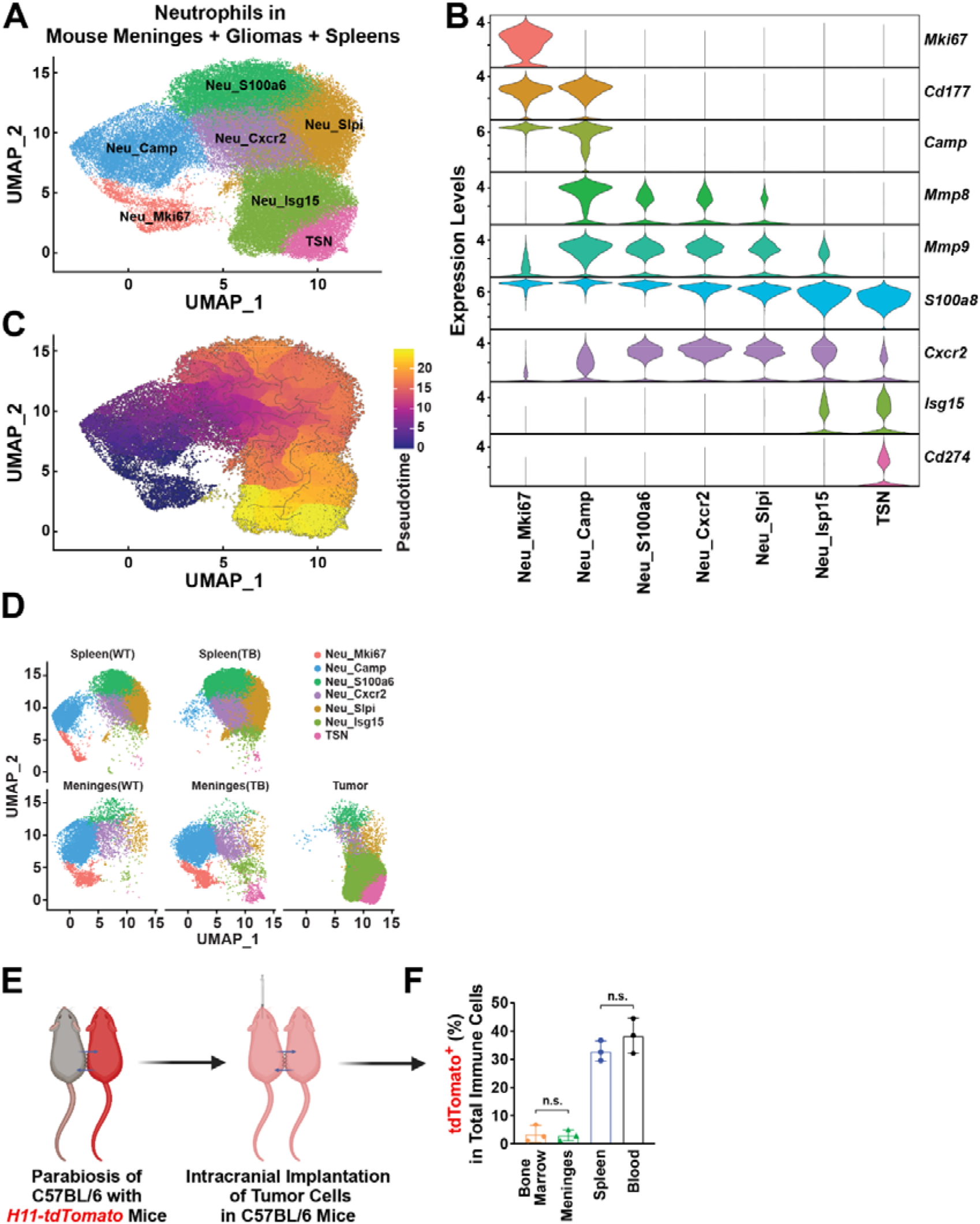
Meninges generates tumor-specific neutrophils in mouse glioma models. Related to Figure 4. **(A to D)** C57BL/6 wild-type mice were intracranially implanted with glioma cells. **(A)** TSNs and other neutrophil subtypes identified in glioma tumors and the meninges and spleens of control or tumor-bearing mice. Neu_Camp, Neu_Cxcr2, Neu_Isg15, Neu_Mki67, Neu_S100a6, Neu_Slpi, neutrophils with corresponding markers; TSN, tumor-specific neutrophils. **(B)** Violin plot of the expression of the proliferation marker gene (*Mki67*), immature neutrophil marker genes (*Cd177* and *Camp*), neutrophil-derived proteolytic enzymes (*Mmp8* and *Mmp9*), neutrophil marker genes (*S100a8* and *Cxcr2*), interferon-stimulated gene (*Isg15*), and immunosuppressive gene (*Cd274*) in neutrophil subtypes. **(C)** Pseudotime trajectory analysis of neutrophil subtypes defined in **(A)**. **(D)** Presence of TSNs and other neutrophil subtypes in the spleens of control or tumor-bearing (TB) mice (upper panels), meninges of control or tumor-bearing mice, and glioma tumors (lower panels). (**E and F**) Parabiosis in mouse glioma models. **(E)** Diagram of the parabiosis of C57BL/6 wild-type mice with *H11-tdTomato* reporter mice followed by the intracranial implantation of glioma cells in C57BL/6 mice. **(F)** The percentage of tdTomato^+^ cells in total immune cells (Cd45^+^) of the bone marrow, meninges, spleens, and blood of C57BL/6 mice after parabiosis. n.s., not significant.

**Supplemental Figure 11.**
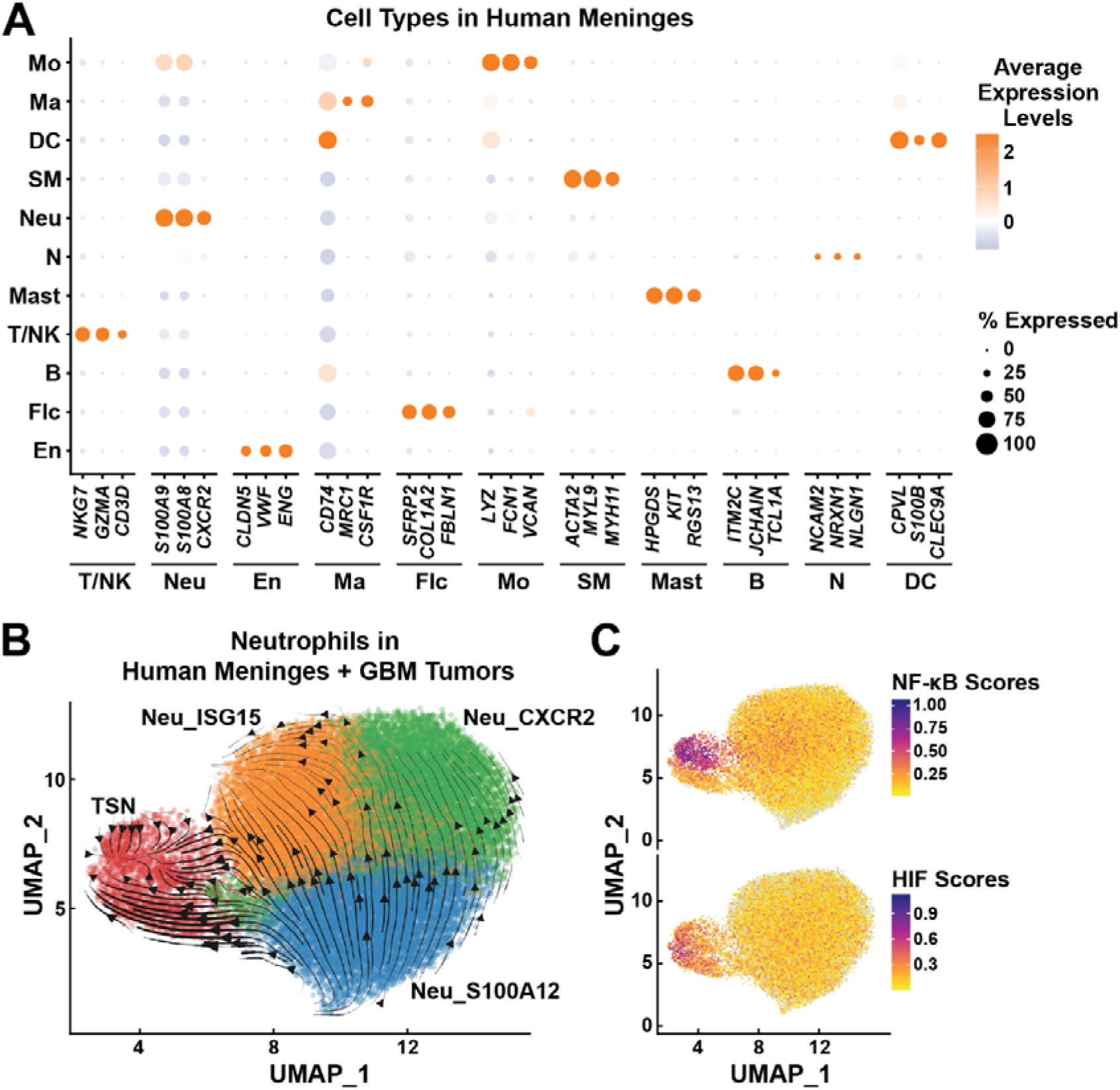
Characterization of meningeal cells of human GBM patients. Related to Figure 1 and Figure 4. **(A)** Dot plot of major cell types identified by profiling the meninges of human GBM patients. Three top marker genes for each cell type are shown. B, B cells; DC, dendritic cells; En, endothelial cells; Flc, fibroblast-like cells; Ma, macrophages; Mast, mast cells; Mo, monocytes; N, neurons; Neu, neutrophils; SM, smooth muscle cells; T/NK, T cells / natural killer cells **(B and C)** Regulation of human TSN development by distinct signaling pathways. **(B)** RNA velocity plot embedded in the UMAP space showing the putative near-transcriptional state of neutrophil subtypes identified in GBM patients’ meninges and tumors. Neu_CXCR2, Neu_ISG15, Neu_S100A12, neutrophils with corresponding markers; TSN, tumor-specific neutrophils. **(C)** Projection of the average expression of 102 NF-κB-target genes (upper panel) and 48 HIF-target genes (lower panel) onto the UMAP plot of neutrophil subtypes.

**Supplemental Figure 12.**
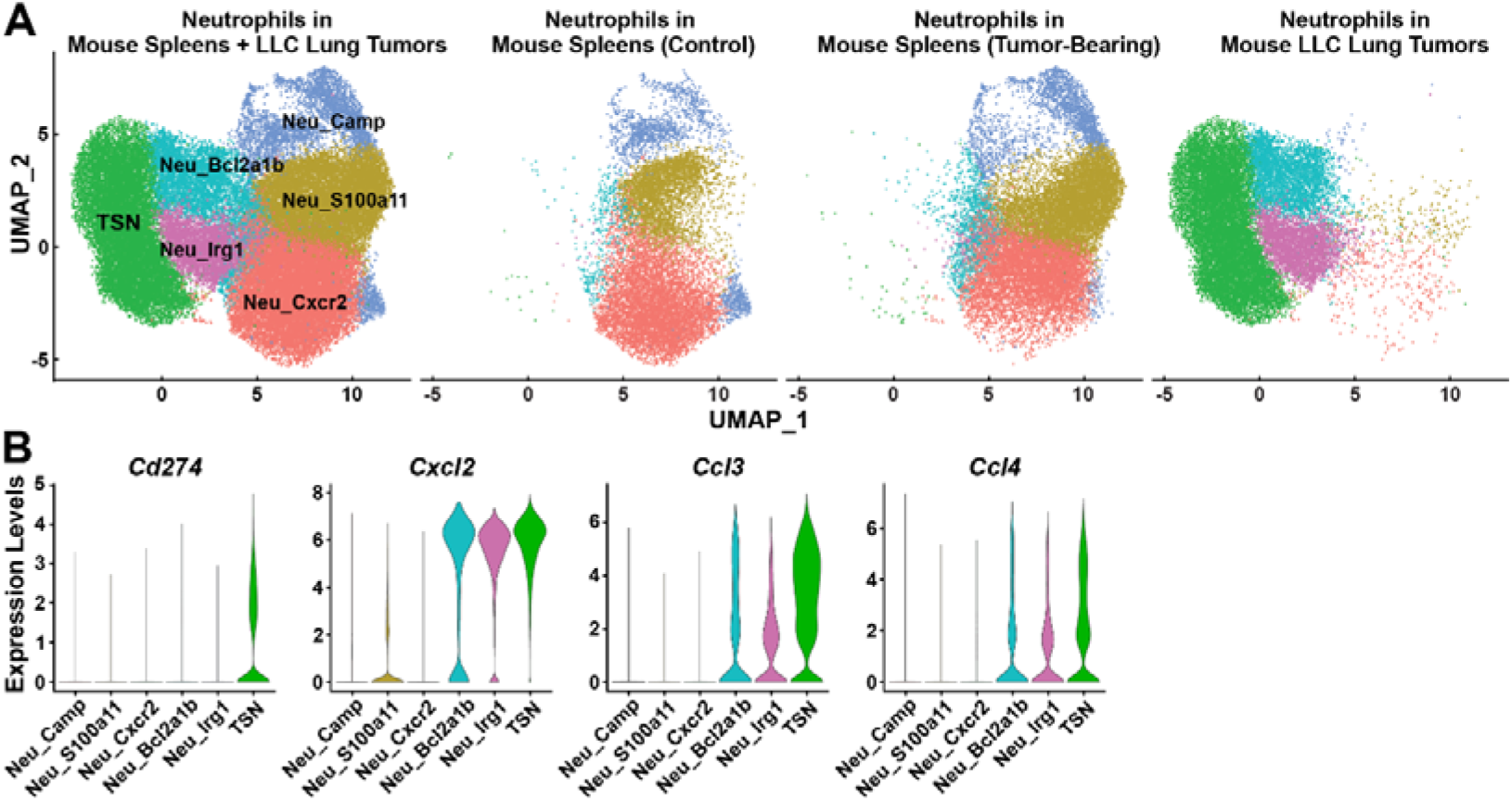
scRNA-seq analysis of neutrophils from mouse spleen and Lewis lung carcinoma (LLC) tumors. **(A)**, UMAP plot illustrating different neutrophil subtypes and neutrophil subtype composition in 3 tissues. **(B)**, Expression of *Cd274*, *Cxcl2*, *Ccl3*, and *Ccl4* in different neutrophil subtypes.

## Materials & Methods

### Human tissue samples and preparations

Human tissue samples were collected in compliance with the protocol approved by the Institutional Ethics Committee of Beijing Tiantan Hospital, Capital Medical University. Informed consent was signed by each involved patient. Detailed patient information is summarized in Table S1. Fresh tissues of human glioblastoma (GBM) or the meninges were immersed in HypoThermosol FRS (BioLife Solutions), and blood samples were collected in lithium heparin tubes. All the human samples were stored at 4°C before further processing.

For the FACS sorting, tissues of human GBM or meninges were briefly rinsed twice in ice-cold phosphate-buffered saline (PBS) before being cut into small pieces. The tissues were digested in RPMI 1640 medium (Thermo Fisher Scientific) containing 0.1 mg/ml Liberase TL (Roche), 20 μg/ml DNase I (Sigma), and 3% heat-inactivated fetal bovine serum (HI-FBS; Sigma) at 37°C for 15 ∼ 30 min, and then mashed through 70-μm cell strainers. The resulting cells were centrifuged at 500 *g* for 5 min and re-suspended in ammonium-chloride-potassium (ACK buffer; Thermo Fisher Scientific) to lyse red blood cells. The cells were centrifuged again at 500 *g* for 5 min and re-suspended in Hanks’ balanced salt solution (HBSS; Thermo Fisher Scientific) containing 3% HI-FBS. The cells were stained by the intended FACS antibodies and sorted on BD FACSAria for single-cell RNA-seq (scRNA-seq) by Singleron. Also, human blood samples were centrifuged at 500 *g* for 5 min, and red blood cells were lysed twice in ACK buffer. The resulting white blood cells were re-suspended in HBSS containing 3% HI-FBS, stained by the intended FACS antibodies, and sorted on BD FACSAria for scRNA-seq by Singleron.

For the FACS analyses, human GBM tissues or meninges were processed as above. The FACS-stained cells were loaded on the BD LSRFortessa, and the data were analyzed by FlowJo (https://www.flowjo.com).

### Mouse information

All the experimental procedures were performed in compliance with the protocols approved by the Institutional Animal Care and Use Committee (IACUC) of the Chinese Institute for Brain Research or Peking University. Mice were maintained on the 12hr/12hr light/dark cycle (light period 7:00 am ∼ 7:00 pm) at ambient temperature (21°C ∼ 24°C) with water and standard chow diet available *ad libitum*.

C57BL/6 wild-type mice and BALB/c nude mice were purchased from Charles River International. *Cxcr2^-/-^* (strain #006848), *Ki67-RFP* (strain #029802), and *E2a-Cre* (strain #003724) mice were from the Jackson Laboratory. *H11-LoxP-ZsGreen-STOP-LoxP-tdTomato* (strain #T006163) mice from GemPharmatech were bred with *E2a-Cre* to generate *H11-tdTomato* mice, which were then used for parabiosis.

### Mouse tumor cell lines

#### LCPN cells

Neural stem cells from the postnatal day 5 (P5) cortex of C57BL/6 wild-type male mice were cultured in the serum-free medium (see below). The cells were sequentially transduced with lentiviruses of the lentiCRISPR-Puro (Addgene plasmid #52961) expressing *Tp53* sgRNA (CCTCGAGCTCCCTCTGAGCC) and the pLKO.1-Hygro (Addgene Plasmid #24150) expressing *Nf1* sgRNA (GTTGTGCTCGGTGCTGACTT). After culture for 2-3 weeks, LCPN cells were obtained for their ability of long-term passage *in vitro*. Notably, due to high expression levels of Y antigens, LCPN cells are tumorigenic only intracranially but not subcutaneously in C57BL/6 female mice.

#### LCPNS cells

The cells isolated from LCPN intracranial tumors of C57BL/6 female mice were cultured under the serum-free condition to obtain LCPNS cells, which spontaneously exhibit lower levels of Y antigens. Of importance, LCPNS cells become tumorigenic subcutaneously in C57BL/6 female mice.

#### LCPNS-SIIN cells

For increasing the tumor antigenicity, LCPNS cells were stably transduced with lentivirus expressing the fusion protein of mScarlet and the MHC class I (H-2Kb) restriction epitope SIINFEKL of ovalbumin (SIIN peptide, Table S4) to obtain LCPNS-SIIN cells. LCPNS-SIIN cells are tumorigenic only intracranially but not subcutaneously in both C57BL/6 male and female mice.

#### LCPNS-SIIN-*Cxcl1*KO cells

LCPNS-SIIN cells were transduced by lentivirus of the pLKO.1-Hygro expressing *Cxcl1* sgRNA1 (ACTTCGGTTTGGGTGCAGTG) and sgRNA2 (GCAGTGGCGAGACCTACCTG).

Single cells were isolated into 96-well plates for clonal selection. Different clones of LCPNS-SIIN-*Cxcl1KO* cells were verified by the DNA sequencing of *Cxcl1* alleles and the ELISA analyses of CXCL1 protein levels in culture media.

#### GL261-SIIN cells

GL261 cell is a gift from Dr. Chao Yan from Nanjing University. GL261 cells were stably transduced with lentivirus expressing the fusion protein of mScarlet and SIIN peptide to obtain GL261-SIIN cells.

#### GL261-SIIN-*Cxcl1* cells

GL261-SIIN cells were stably transduced with a lentivirus expressing mouse *Cxcl1*. GL261-SIIN-*Cxcl1* cells were verified by the ELISA analyses of CXCL1 protein levels in culture media.

Mouse neural stem cells, LCPN cells, LCPNS cells, LCPNS-SIIN cells, and LCPNS-SIIN-*Cxcl1*KO cells were cultured at 37°C under 5% CO_2_ / 5% O_2_ in Dulbecco’s Modified Eagle Medium/Nutrient Mixture F-12 (DMEM/F12 medium, Gibco) supplemented with 1% N2 (Gibco), 1% B-27 (Gibco), 1% GlutaMAX (Gibco), 100 U/ml penicillin, 100 μg/ml streptomycin, 16.6 mM *D*-glucose, 5mM HEPES, 50 nM 2-mercaptoethanol (Sigma), 30 ng/ml EGF (Novoprotein), and 30 ng/ml FGF2 (Origene).

GL261 cells, GL261-SIIN cells, and GL261-SIIN-*Cxcl1* cells were cultured at 37°C under 5% CO_2_ in DMEM medium (Gibco) supplemented with 10% FBS (VISTECH), 100 U/ml penicillin, and 100 μg/ml streptomycin.

### Mouse tumor implantation

For the subcutaneous implantation of glioma cells, 5 × 10^5^ LCPN cells, LCPNS cells, or LCPNS-SIIN cells embedded in 100 μl Matrigel (Corning) were injected into the right flank of 6 ∼ 8 week-old female mice. Tumor dimensions were measured every other day with a caliper, and tumor volume was calculated as volume = 0.5 × length × width × width.

For the intracranial implantation of glioma cells, 6 ∼ 8 week-old female mice were anesthetized and head-fixed on the stereotactic frame. The head skin was shaved and prepared with iodine and 75% alcohol. A skin incision was made along the midline to expose the skull. The following coordinates for injection were measured relative to the bregma: anteroposterior = 0.5 mm, medial-lateral = 1.9 mm, and dorsal-ventral = −3.2 mm. A small hole was drilled through the skull at the target site, through which a Hamilton syringe needle (701N, Hamilton) was inserted into the brain. 5 × 10^5^ LCPN cells, LCPNS cells, or LCPNS-SIIN cells embedded in 5 μl

Matrigel (Corning) was delivered at the injection rate of 1.5 μl/min. The needle was kept in place for an additional 5 min to prevent the leakage of injected cells. Bone wax was used to cover the small hole on the skull to avoid tumor cells overflowinbg.

The mice were monitored for body weight every other day and euthanized when a 25% drop in body weight was observed.

### Mouse parabiosis

Each pair of mice was housed together for 1 week before the surgery. The mice were anesthetized, and the skin on one side of each mouse was shaved and prepared with iodine and 75% alcohol. A longitudinal incision was made along the side of each mouse, and the skin was carefully separated from the underlying connective tissues. A longitudinal incision of approximately 10 mm was then made on the exposed peritoneum of each mouse. The incision sites of the two mice were sutured together to establish the connection of vascular systems. In addition, the scapulae on the incision sides of the mice were sutured together to help hold the parabiotic pair. The skin incisions of the mice were closed together by surgical staples. Each parabiotic pair was housed in a cage and utilized for experiments at 18 days post-surgery.

### FACS analyses of mouse tissues

Mouse glioma tumors were freshly dissected and cut into small pieces on ice. The tissues were digested in an adequate volume of Accutase (BioLegend) at 37°C for 15 min and then mashed through 70-μm cell strainers. Mouse meninges were freshly dissected and digested in RPMI 1640 medium containing 0.1 mg/ml Liberase TL, 20 μg/ml DNase I, and 3% HI-FBS at 37°C for 15 min. The meningeal tissues were then mashed through 70-μm cell strainers. Mouse spleens were freshly dissected and directly mashed through 70-μm cell strainers in HBSS containing 3% HI-FBS. Mouse bone marrow was flushed out from the femurs with PBS, and the cells were filtered through 70-μm cell strainers. Mouse peripheral blood samples were collected via retro-orbital bleeding into PBS containing 5 mM EDTA. The resulting cell suspensions of mouse tissues were centrifuged at 500 *g* for 5 min and re-suspended in ACK buffer to lyse red blood cells. The cells were centrifuged again at 500 *g* for 5 min, re-suspended in HBSS containing 3% HI-FBS, and stained by the intended FACS antibodies. The stained cells were sorted on BD FACSAria for scRNA-seq by Singleron. Alternatively, the stained cells were processed on BD LSRFortessa, and the data were analyzed by FlowJo (https://www.flowjo.com).

### Histochemistry or immunohistochemistry staining

For the H&E staining, formalin-fixed human gliomas or mouse brains were embedded in paraffin. 5 μm-thick sections were cut on a microtome and stained with the Hematoxylin and Eosin Staining Kit (Solarbio).

For the immunohistochemistry staining, 5 μm-thick paraffin sections were deparaffinized and stained with anti-human KI67 (1:500, TA802544, Origene), anti-mouse Ki67 (1:800, #12202, Cell Signaling Technology), anti-mouse Ly6G (1:200, #87048, Cell Signaling Technology), anti-mouse Cd8a (1:800, #98941, Cell Signaling Technology), or anti-mouse Cd274 (1:200, #64988, Cell Signaling Technology). The sections were incubated with the anti-mouse/rabbit HRP secondary antibody and detected by DAB solution, according to the manual of the Two-step Universal Kit (ZSGB-BIO). Histological images were analyzed using HALO (Indica Labs-Multiplex IHC v3.1.4).

For the immunofluorescence staining, 5 μm-thick paraffin sections were deparaffinized and stained with anti-human CD66b (1:100, GTX19779, GeneTex), anti-human CD274 (1:100, #13684, Cell Signaling Technology), anti-mouse Ly6G (1:300, GB11229, ServiceBio), or anti-mouse Cd274 (15μg/mL, MAB1561, R&D Systems). The sections were further stained with Alexa Fluor-conjugated secondary antibodies (Thermo Fisher Scientific) and mounted with DAPI. The images were obtained by epifluorescence microscopy.

### Quantitative PCR analyses

Total RNAs of mouse tissues or cultured glioma cells were extracted by the FastPure Cell/Tissue Total RNA Isolation Kit (Vazyme), reverse transcribed into cDNAs by the HiScript III All-in-one RT SuperMix Perfect (Vazyme), and analyzed by the SYBR Green Real-Time PCR Kit (Thermo Fisher Scientific). The primer sequences used for quantitative PCR are listed in Table S4.

### scRNA-seq data preprocessing

Data preprocessing was performed according to the manufacturer’s reference instructions. For the scRNA-seq results using 10X genomics products, we followed the CellRanger website (https://support.10xgenomics.com/single-cell-gene-expression/software/pipelines/late st/ advanced/references), using CellRanger (v7.0.1) for sequence alignment and gene expression quantification. For the scRNA-seq results of Singleron products, we followed the CeleScope instructions (https://github.com/singleron-RD/CeleScope), using CeleScope (v1.13.0) for sequence alignment and gene expression quantification. The reference genome version used for sequence alignment was GRch38 for the human and mm10 for the mouse. We filtered for genes expressed in only 10 or fewer cells, as well as cells expressing gene number <200, mitochondrial reads >10%, and ribosome genes proportioned >20%. We used the Solo^30^ tool in scvi-tools to remove doublet using the first 2000 top variant genes and default parameters.

### Dimensionality reduction, clustering, and cell type determination of scRNA-seq

We used single-cell Variational Inference (scVI)^31^ from scvi-tools according to software instructions (https://docs.scvi-tools.org/en/stable/tutorials/index.html) to integrate different samples of scRNA-seq results together. We used scanpy^32^ to cluster the integrated data, and the resulting H5ad files were converted into Seurat objects using Sceasy (https://github.com/cellgeni/sceasy) and run through Rstudio with R v4.2.2^33^, using Seurat v.4’s^34^ FindAllMarkers to determine and visualize the markers for each cluster. We used a combination of ScType^35^ and PanglaoDB^36^ for the initial determination of cell types. For macrophage^37^ and neutrophil subtypes^38^, we referenced the published results for further determination.

### RNA velocity analysis of scRNA-seq

We used the default parameters of scVelo^39^ to calculate RNA velocity according to software instructions (https://smorabit.github.io/tutorials/8_velocyto/). scVelo’s input was calculated using the Python version of velocyto. The pseudotime trajectory analysis was done by monocle3^40^ (https://cole-trapnell-lab.github.io/monocle3/docs/getting_started/) using default parameters. The modulation network was explored using the docker version of pySCENIC (0.12.0)^26^ (https://hub.docker.com/r/aertslab/pyscenic). Factors differentially expressed in different cell types were identified based on AUC scores using FindAllMarker, which were visualized using ComplexHeatmap^41^ and Clusterprofiler^42^. Sceasy was used to complete the conversion of H5ad and Seurat objects.

### Bulk RNA-seq analyses

STAR 2.7.3a^43^ was used to align the short sequence sequencing files with the reference human or mouse genomes, which was obtained from Ensembl (v. 98). The FeatureCount^44^ program in the Subread package^45^ was used to quantify gene expression. Gene expression differential analysis was carried out by edgeR^46^. Signaling pathway enrichment was done by the GSEApreranked algorithm in GSEA v4.2.3^47^ using gene differential expression results as input.

TCGA and CGGA databases^21^ were obtained from GDAC Firehose of Broad Institute (https://gdac.broadinstitute.org) and Beijing Neurosurgical Institute (http://www.cgga.org.cn/), respectively. was used to calculate the TSN score in each sample. The top 1/3 tumor samples with the highest TSN scores and the bottom 1/3 tumor samples with the lowest TSN scores in the same dataset were used to compare the survival differences of patients.

### Statistical methods

Student’s *t*-tests (two-tailed unpaired), ANOVA with *post hoc* tests, or log-rank tests were performed using GraphPad Prism 8.4.3 (http://www.graphpad.com/scientific-software/prism). Statistical details of the experiments are included in the figure legends.

